# The Cyclic AMP Receptor Protein Regulates Quorum Sensing and Global Gene Expression in *Yersinia pestis* During Planktonic Growth and Growth in Biofilms

**DOI:** 10.1101/742031

**Authors:** Jeremy T. Ritzert, George Minasov, Ryan Embry, Matthew J. Schipma, Karla J. F. Satchell

**Author notes:** To whom correspondence should be addressed: Karla J. F. Satchell. Department of Biology, Loyola University Chicago, Chicago, IL, 60660, USA.

## Abstract

Cyclic adenosine monophosphate (cAMP) receptor protein (Crp) is an important transcriptional regulator of *Yersinia pestis.* Expression of *crp* increases during pneumonic plague as the pathogen depletes glucose and forms large biofilms within lungs. To better understand control of *Y. pestis* Crp, we determined a 1.8 Å crystal structure of the protein-cAMP complex. We found that compared to *Escherichia coli* Crp, C helix amino acid substitutions in *Y. pestis* Crp did not impact cAMP dependency of Crp to bind DNA promoters. To investigate *Y. pestis* Crp-regulated genes during plague pneumonia, we performed RNA-sequencing on both wild-type and Δ*crp* mutant bacteria growing in planktonic and biofilm states in minimal media with glucose or glycerol. *Y. pestis* Crp is found to dramatically alter expression of hundreds of genes dependent upon carbon source and growth state. Gel shift assays confirmed direct regulation of the *malT* and *ptsG* promoters and Crp was then linked to *Y. pestis* growth on maltose as a sole carbon source. Iron-regulation genes *ybtA* and *fyuA* were found to be indirectly regulated by Crp. A new connection between carbon source and quorum sensing was revealed as Crp was found to regulate production of acyl-homoserine lactones (AHLs) through direct and indirect regulation of genes for AHL synthetases and receptors. AHLs were subsequently identified in the lungs of *Y. pestis* infected mice when *crp* expression is highest in *Y. pestis* biofilms. Thus, in addition to well-studied *pla,* other Crp-regulated genes likely have important functions during plague infection.

**IMPORTANCE:** Bacterial pathogens have evolved extensive signaling pathways to translate environmental signals into changes in gene expression. While Crp has long been appreciated for its role in regulating metabolism of carbon sources in many bacterial species, transcriptional profiling has revealed that this protein regulates many other aspects of bacterial physiology. The plague pathogen, *Y. pestis,* requires this global regulator to survive in blood, skin, and lungs. During disease progression, this organism adapts to changes within these niches. In addition to regulating genes for metabolism of non-glucose sugars, we find the Crp regulates genes for virulence, metal acquisition and quorum sensing by direct or indirect mechanisms. Thus, this single transcriptional regulator, that responds to changes in available carbon sources, can regulate multiple critical behaviors for causing disease.

## INTRODUCTION

*Yersinia pestis* is the etiological agent of pneumonic, bubonic and septicemic plague. While cases of plague are rare, *Y. pestis* evolved to adapt to environmental changes between its life cycle in fleas, rodents and mammals, including humans (1). The flea, bubo (lymph nodes), blood, and lungs differ in temperature, nutrients, and defense systems. To interpret changes in these environments, *Y. pestis* encodes more than two dozen two-component systems (2), three quorum sensing systems (3), and additional transcriptional regulators with specialized functions. This sensor network allows *Y. pestis* to translate changes in its extracellular environment into altered gene expression to promote growth and pathogenesis (4, 5).

The 3’,5’-cyclic adenosine monophosphate (cAMP) receptor protein (Crp) is required for *Y. pestis* virulence in mice (6). Crp is a 23.5 kDa transcriptional regulator that forms a dimer after binding its ligand cAMP (7). Dimerization allows for activation or repression of gene expression by binding to the promoter region of its target genes (8). The activity of adenylate cyclase (CyaA) regulates intracellular concentrations of cAMP *via* the conversion of ATP into cAMP. This activity is increased at 37°C compared to lower temperatures, suggesting cAMP-Crp signaling is important during mammalian infection (9, 10). In addition, phosphorylated EIIA from the phosphotransferase system activates CyaA when glucose is absent (11). During catabolite repression, the transport of glucose through PtsG depletes intracellular concentrations of phosphorylated EIIA as the phosphate group is transferred to the incoming glucose molecule.

While known for its role during catabolite repression and regulation of the *lac* operon, it is now appreciated that Crp regulates expression of other genes, including factors important during infection to connect changes in glucose availability to regulation of bacterial behaviors. Across γ-proteobacteria, Crp regulates biofilm formation (12, 13), capsule production (14), the DNA damage response (15), toxin production (16, 17), luminescence (18), and iron acquisition (19).

Other proteins can regulate expression of the *crp* gene, suggesting that multiple input signals modulate expression of the Crp regulon. The two-component system PhoPQ and the small RNA chaperone Hfq regulate *crp* in *Y. pestis* (6, 20, 21). In turn, Crp regulates expression of the type III secretion system and the Pla protease essential for pneumonic plague (22–24). Crp also promotes biofilm formation *via* a mechanism involving the RNA-binding regulatory protein CsrA (13). Production of the main constituent of biofilms, poly-*N*-acetylglucosamine, is increased at 37°C in fully virulent strains of *Y. pestis,* suggesting additional roles for biofilm formation during mammalian infection (25). Further, we recently demonstrated increased expression of *crp* within biofilms in the lungs of *Y. pestis*-infected mice, revealing a link between *in vivo* biofilms, glucose availability, and essential virulence gene expression (26).

In this study, we utilize global transcriptional profiling between planktonic and biofilm states in the presence of glucose and glycerol to reveal previously unrecognized Crp-regulated genes in *Y. pestis.* We find that Crp indirectly represses genes required for siderophore biosynthesis and stimulates genes for carbohydrate uptake and metabolism, particularly the use of maltose as an alternative carbon source. Unexpectedly, Crp was found to promote expression of the acyl-homoserine lactone (AHL) quorum sensing genes. Crp directly binds to the promoter for the AHL-receptor, *ypeR,* and thereby controls efficient production of AHLs within biofilms.

## RESULTS AND DISCUSSION

### Experimental setup

Pneumonic plague is a biphasic disease consisting of an early non-inflammatory phase and a damaging pro-inflammatory phase (27). As pneumonia develops, the lungs fill with *Y. pestis,* neutrophils, and fluid (Fig. 1A). *Y. pestis* proliferates to ∼10^9^ colony forming units (CFUs), forms large biofilms in the lungs, and consumes all available glucose in the process. The declining concentration of glucose activates expression of *crp* within biofilms (26). Expression of the Crp-activated gene *pla,* which is required for *Y. pestis* to grow within the lungs and disseminate to other organs, also increases (22). To better understand the changes in the lung environment, we sought to identify additional Crp-regulated genes dependent upon growth in biofilms and glucose-limiting conditions.

**FIG. 1.**
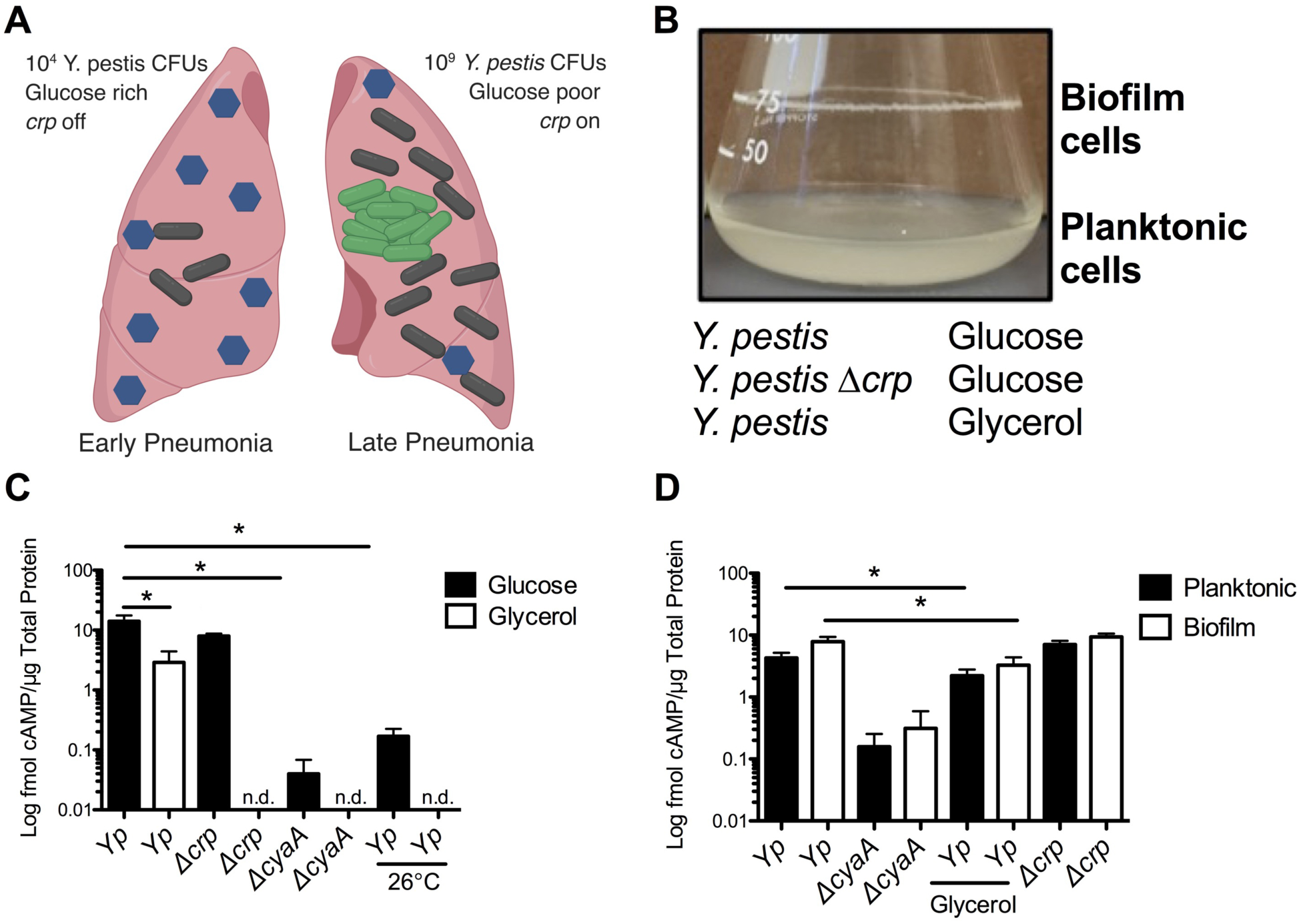
Experimental set up for RNA-seq. (A) Diagram of changes during progression of pneumonic plague. Blue hexagons represent glucose. Black and green bacilli represent *Y. pestis* and *Y. pestis* expressing *crp* in biofilms, respectively. (B) 10 mL culture of *Y. pestis* grown in TMH for 18 hours at 37°C showing the biofilm formed at the air-liquid interface. RNA was isolated from *Y. pestis* or *Y. pestis Δcrp* cultures grown in TMH with 0.2% glucose or glycerol under these conditions in triplicate. (C) Intracellular concentrations of cAMP in *Y. pestis (Yp)*, *Y. pestis Δcrp*, or *Y. pestis ΔcyaA* grown for six hours in TMH with 0.2% glucose (black bars) or glycerol (white bars) at 37°C or 26°C as indicated. n.d. = not done. (D) Intracellular concentrations of cAMP in *Y. pestis (Yp)*, *Y. pestis Δcrp*, or *Y. pestis ΔcyaA* grown for 18 hours in TMH with 0.2% glucose or glycerol in planktonic (black bars) or biofilm (white bars) states at 37°C. Data represent mean and SEM from three independent experiments. \**p*<0.05 and \*\**p*<0.01 from Student’s *t* test.

To recapitulate human infection conditions, *Y. pestis* was grown at 37°C with shaking in defined liquid culture media with glucose to mimic early pneumonia or with glycerol to mimic later infection after glucose is consumed. As the cultures were aerated by shaking, *Y. pestis* formed a biofilm at the air-liquid interface, thereby facilitating comparison of gene expression between the planktonic state representing early infection and the biofilm state of later infection (Fig. 1B). In addition, a Δ*crp* mutant was included in parallel to facilitate identification specifically of Crp-regulated genes under these conditions. These combined six experimental conditions allow for identifying glucose-, Crp-, and biofilm-dependent genes that may play a role during pneumonic plague infection.

### Carbon source and growth state do not affect cAMP requirement for Crp binding to DNA

A potential concern of this experimental set up is that differences in expression of Crp-regulated genes could be affected by changes in the activity of CyaA and the phosphotransferase system (28), leading to differences in Crp activity rather than expression of *crp*. To control for this possibility, we measured cAMP concentrations in *Y. pestis* under all experimental conditions. In contrast to an expected increase, we observed a decrease in cAMP concentrations in *Y. pestis* grown in glycerol (Fig. 1C). This was also reported in *Vibrio fischeri* in which intracellular cAMP was reduced, but total cAMP concentration (including extracellular cAMP) was higher (29). Deletion of the adenylate cyclase gene *cyaA* reduced cAMP levels, while deletion of *crp* had no effect. cAMP concentrations were much higher in cells grown at 37°C *vs*. 26°C, suggesting the importance of Crp regulation during mammalian infection. cAMP concentrations were not significantly affected between planktonic-and biofilm-grown cells (Fig. 1D). These data indicate that any observed changes in expression of Crp-regulated genes by RNA-sequencing (RNA-seq) would not be due to differences in the cAMP levels.

Another potential caveat of this experimental set up is the possibility that Crp could bind and regulate genes in *Y. pestis* entirely independently of cAMP. Earlier mutational studies on *E. coli* showed that the specific double mutant variant T128L/S129I had extremely high cAMP-independent DNA binding affinity, comparable with the activity of cAMP-bound wild-type Crp (30). *Y. pestis* shares 99% sequence homology to Crp from *E. coli* with differences located on the C helix at positions 119, 123 and 127. We considered that these sequence difference could affect cAMP binding or dimerization and thus cause changes in DNA binding. For this we carried out structural studies using protein crystallography and solved the crystal structure of *Y. pestis* Crp in complex with cAMP. Crp was crystallized as a dimer and each monomer had cAMP bound to the binding site (Fig. 2A). We compared our structure with the structure of *E. coli* Crp bound to cAMP (PDB ID 4R8H, (30)). The structures closely aligned with r.m.s.d of 0.8 Å for 185 pruned C*α* atom pairs and 1.1 Å for all 200 pairs (Fig. 2B). Notably, in the *Y. pestis* structure, C helix residues S119, N123, and I127 (unlike S129 in the constitutive binding mutant of *E. coli* Crp) are directed away from cAMP suggesting these altered amino acids should not impact cAMP binding (Fig. 2C). The side chain of residue N123 is involved in the dimer formation and directly interacts with the side chain of E78 from another chain. The N123 exchange for arginine, which occurs in *Y. pestis,* is disrupting this interaction, but the S119 replacement for alanine restores the hydrogen bond to E78 and thus these two mutations do not alter the stability of dimer. The V127 in Crp from *E. coli* is pointed away from dimerization interface and it is not involved in the cAMP binding, so that isoleucine at this position in *Y. pestis* does not have any impact contribute to binding of cAMP or dimer formation. As confirmation of our findings, an electrophoretic mobility shift assay (EMSA) in the presence of cAMP (see Fig. 5A) indicates that Crp requires cAMP to bind a DNA fragment that corresponds to the *pla* promoter. This cAMP-dependence is also consistent with studies of Crp from other *Y. pestis* isolates and from *Yersinia pseudotuberculosis* (23, 31, 32).

**FIG. 2.**
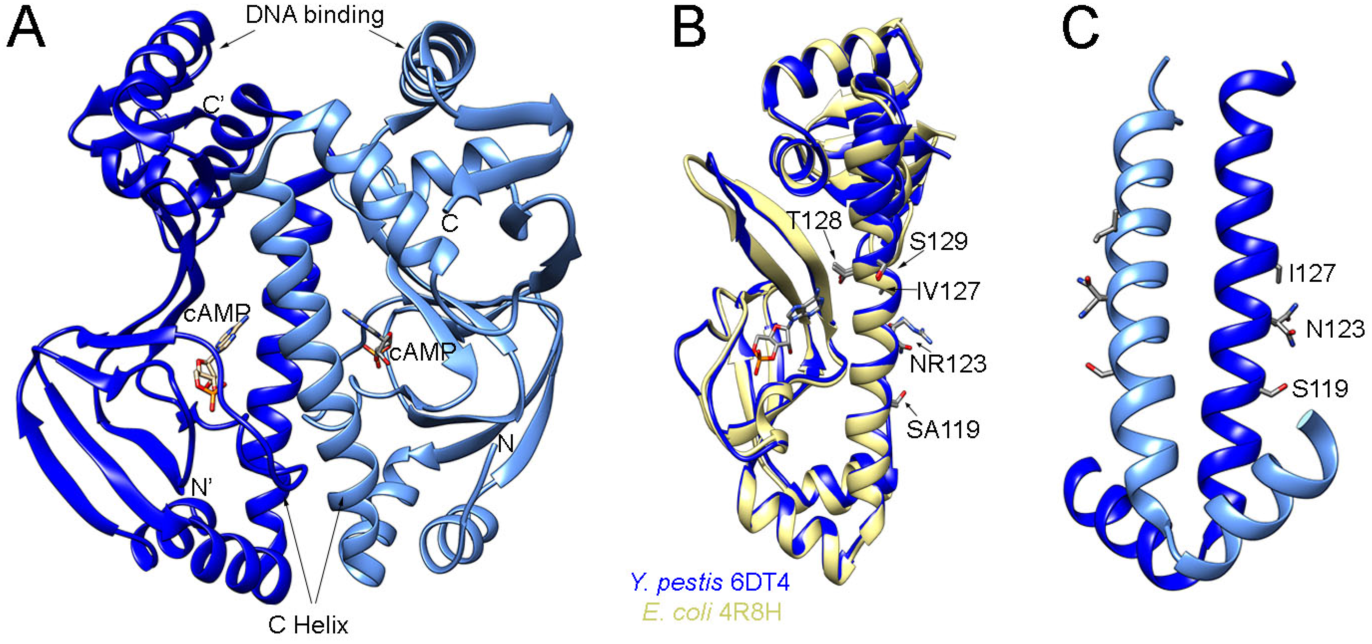
Structure of *Y. pestis* Crp closely aligns with *E. coli* Crp. (A) 1.8 Å resolution structure of *Y. pestis* Crp dimer bound to cAMP (PDB ID 6DT4). (B) Overlap of *Y. pestis* Crp chain A (blue) with *E. coli* Crp (yellow) (PDB ID 4R8H, chain A, (30)). Residues that differ between *Y. pestis* and *E. coli* are marked with *Y. pestis* residue indicated first. Residues modified in *E. coli* that result in constitutive binding to DNA are also marked. (C) Dimer interface of *Y. pestis* Crp showing residues that differ from *E. coli*.

Thus, cAMP concentration data, structural comparison, and EMSA data suggest that Crp function is controlled by cAMP in *Y. pestis*, but that cAMP concentrations do not vary dramatically in the conditions surveyed in this study. Thus, we expect that variations in gene expression noted in this study will be linked specifically to Crp. This suggestion is consistent with our prior finding that regulation of *pla* (between planktonic and biofilm-grown cells) depends on differential expression of Crp as opposed to changes in cAMP (26).

### Global analysis of gene expression

We subsequently performed RNA-seq on total RNA isolated from *Y. pestis* pCD1^-^ and the Δ*crp* mutant under all experimental conditions. RNA from *Y. pestis Δcrp* growing planktonically and in biofilms was collected to identify Crp-regulated genes. Deep sequencing reads were mapped to the *Y. pestis* chromosome and plasmids pPCP1 and pMT1. Reads were also mapped to the location of annotated sRNAs in *Y. pestis* (33). The raw RNA-seq datasets were deposited into GenBank (GSE135228). A cut-off for significantly differentially expressed genes was set at log_2_ fold-change >1 or <-1 with a false-discovery *p*-value <0.05. A complete list of differentially expressed genes can be found in Datasets S1 and S2. Principal component analysis revealed close association of most of the replicates across all conditions (Fig. S1). Crp and carbon source altered expression of thousands of *Y. pestis* genes in planktonic and biofilm growth states (Figs. 3A-D). Indeed, 1,200 unique protein-coding genes were impacted by the presence of *crp* (713 Crp-activated and 487 Crp-repressed), while the presence of glucose altered expression of 1,872 unique genes between planktonic and biofilm growth states.

Only 47 genes were differentially expressed strictly between planktonic- and biofilm-grown cells (Fig. 3F). The low number of differentially expressed genes could be due to overlap of planktonic cells with biofilm cells at the air-liquid interface or heterogeneous expression of genes throughout the biofilm. Genes required for biofilm formation such as diguanylate cyclases, *hmsT* and *ypo0449*, were not differentially expressed between planktonic and biofilm cells (12, 34, 35). A similar observation was reported by Vadyvaloo et al. wherein these genes were also not significantly different between planktonic cells and biofilms in flow cells (36). Genes previously known to influence biofilm formation namely, *crp* (12), *csrA* (13), *rcsAB* (37), *rovM* and *rovA* (38), *phoPQ* (39), *fur* (40), and *hfq* (35), were also not differentially expressed. These data more likely suggest genes for regulating biofilm formation are controlled by environmental factors, such as temperature and growth in the flea (41, 42). Carbon source is also known to play a role (13), and while we observed no difference in expression of the diguanylate cyclases in biofilms, they were significantly upregulated in glycerol compared to glucose (Dataset 1).

**FIG. 3.**
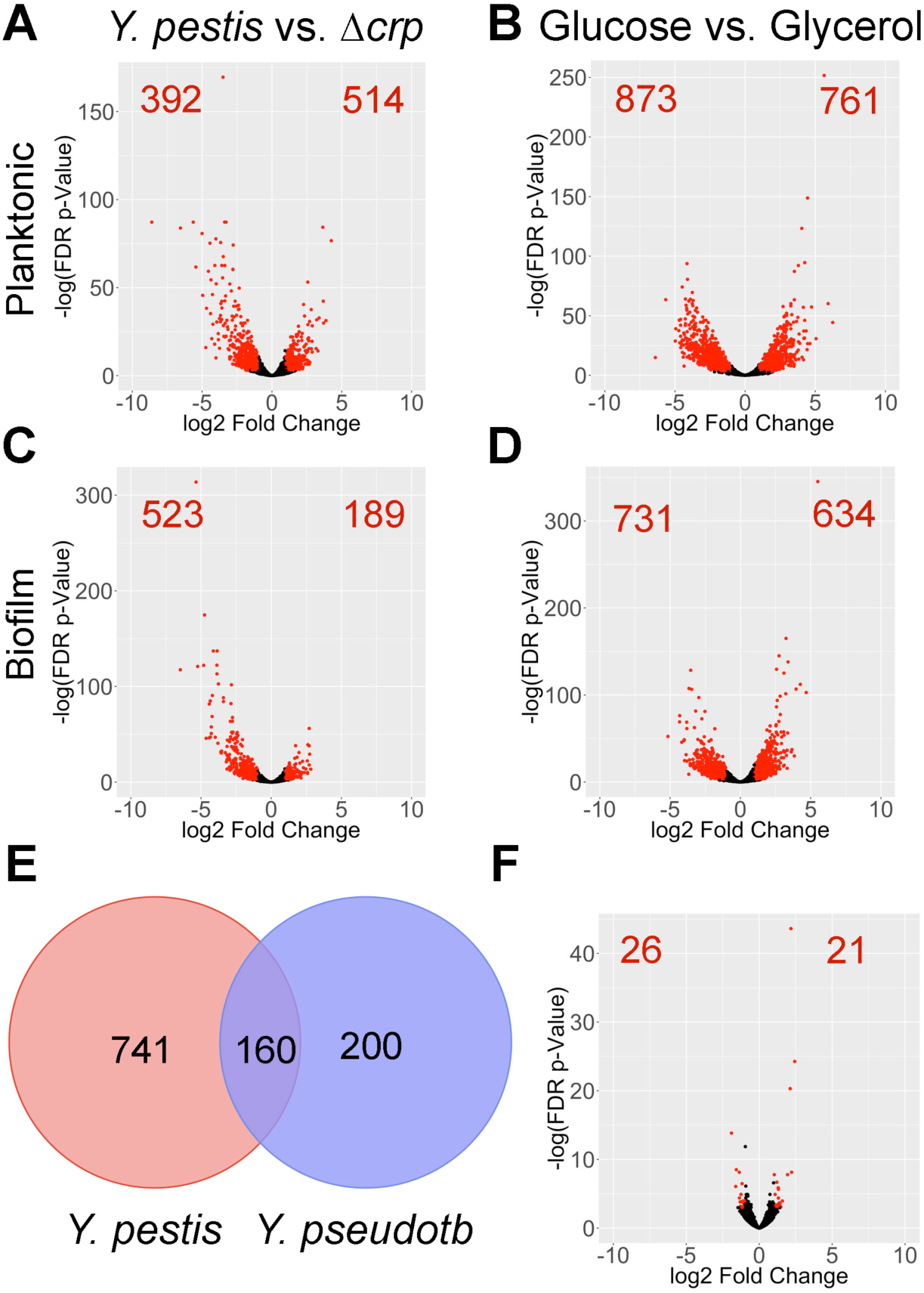
Identification of Crp-regulated, Biofilm-regulated, and Glucose-regulated genes in *Y. pestis.* Volcano plots of differentially expressed genes between *Y. pestis* vs. *Y. pestis* Δ*crp* and *Y. pestis* grown in 0.2% glucose vs. glycerol in (A-B) planktonic and (C-D) biofilm states. Each individual dot represents a gene of *Y. pestis*, red dots and numbers indicate genes with log2 fold change >1 or <-1 and a false discovery rate *p* value <0.05. (E) Venn diagram of Crp-regulated genes in *Y. pestis* from this study and Crp-regulated genes in related *Y. pseudotuberculosis* YPIII (31). (F) Volcano plot of differentially expressed genes between *Y. pestis* grown in planktonic and biofilm state in the presence of 0.2% glucose.

**FIG. 4.**
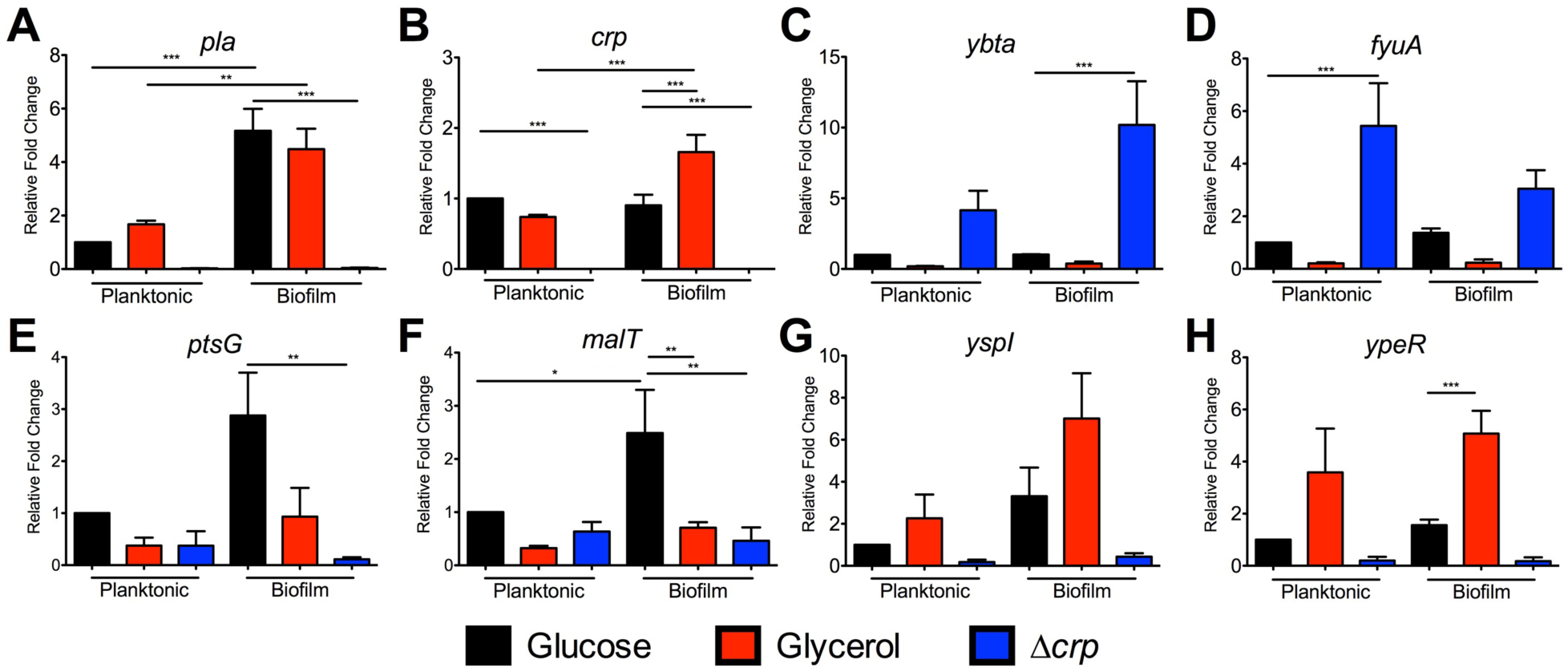
qRT-PCR of Crp-regulated genes identified from RNA-seq. RNA was isolated from cultures grown under the same conditions for RNA-seq. qRT-PCR for (A) *pla*, (B) *crp*, (C) *ybtA*, (D) *fyuA*, (E) *ptsG*, (F) *malT*, (G) *yspI*, and (H) *ypeR*. Data represent the mean and SEM from four independent experiments. \**p*<0.05, \*\**p*<0.01, \*\*\**p*<0.001 from one-way ANOVA with Bonferroni’s multiple comparisons test.

Significant genes were categorized and enriched by biological process with GeneOntology at geneontology.org using *Yersinia pestis* as a reference list (Dataset 3)(43, 44). Categorizing Crp-activated genes in planktonic or biofilm cells returned an enrichment for DNA-templated transcriptional regulators involved in a wide range of biological processes including catabolism of carbohydrates (*malT* and *araC*), amino acid metabolism (*leuO*), and quorum sensing (*ypeR* and *yspR*). This category was also enriched in glucose-repressed genes (i.e. glycerol-activated) with overlap of multiple transcriptional regulators including *malT* and *ypeR.* We anticipated overlap between Crp-activated and glucose-repressed genes, as well as between Crp-repressed and glucose-activated genes based on the function of cAMP-Crp. By filtering these datasets for genes increased in glycerol compared to glucose and increased in *Y. pestis* compared to Δ*crp,* we could better identify cAMP-Crp-activated (or repressed) genes (Table S2). Importantly, the *pla* gene, known to be Crp-activated and glucose-repressed and active during the end stages of pneumonic plague (6, 23), filtered into the Crp-activated category. Several Crp-regulated genes from a previous microarray of *Y. pestis* (*pim, pst, ptsG, araF, rpoH, yfiA* and *ompC*)(32) were also identified. While Gene Ontology identified the quorum sensing transcriptional regulators, *yspR* and *ypeR,* as Crp-activated, the corresponding acyl-homoserine lactone synthetases (*yspI* and *ypeI*) also were identified as CRP-activated (Table S2).

Furthermore, while we observed few differentially expressed genes when comparing *Y. pestis* in biofilms vs. *Y. pestis* planktonic cells, genes and enriched pathways classified as activated or repressed by Crp or glucose differed between planktonic and biofilm cells. In other words, which genes or pathways are turned on by Crp or respond to changes in carbon source depend on whether *Y. pestis* is growing in a planktonic or biofilm state. We subsequently focused on identifying Crp-regulated genes that may play a role during the progression of pneumonic plague as *Y. pestis* forms biofilms and the environment switches from Crp-repressive to Crp-active.

We also observed overlap between Crp-regulated genes identified here for *Y. pestis* and previously published expression profiling for *Y. pseudotuberculosis* compared to its isogenic Δ*crp* mutant (45). All total, 160 genes were shared across these datasets as differentially expressed dependent upon Crp, despite differing culture conditions and cutoffs for significance (Fig. 3E). An additional five genes (*rseC, rpsL, rpsG, rpmA,* and *ybiT*) were Crp-activated in one species, but Crp-repressed in the other. Differences between the two datasets may also result from how the *crp* gene is regulated amongst the two species. The small RNA chaperone, Hfq, is required for full production of Crp in *Y. pestis,* but not in *Y. pseudotuberculosis* (6, 31). The PhoP response regulator is an activator of *crp* expression in one strain of *Y. pestis* (21), but variation in the DNA-binding domain of PhoP between *Yersinia* strains alters transcription of its target genes (46). In addition, these differences in regulation of the *crp* gene and the extent to what genes are regulated by Crp likely result from changes in the promoter region of genes and reflect adaptations to the different environments *Y. pestis* and *Y. pseudotuberculosis* inhabit.

### Crp indirectly represses expression of genes for Yersiniabactin biosynthesis and uptake

Among all the genes identified as controlled by Crp in the RNA-seq dataset, it was noted that Crp represses expression of genes for metal acquisition (Dataset S1). These included genes for biosynthesis and uptake of the iron siderophore Yersiniabactin (Ybt), *fyuA,* the gene for the Ybt receptor, and *ybtA*, the gene for the transcriptional regulator that controls expression of the Ybt locus (Table 1) (47). Consequently, expression of the *irp* genes, required for Ybt production, and genes for heme transport were also decreased, but genes for the Yfe and Feo transport systems were not significantly differentially expressed (Table 1). Crp also represses expression of *ybtX* (*irp8* in strain CO92), a known virulence factor and trigger of inflammation during pneumonic plague (48, 49).

qRT-PCR supported the RNA-seq results as transcript levels of *ybtA* and *fyuA* were reduced in glycerol-grown cultures and also increased in the Δ*crp* mutant (Figs. 4C-D). Crp-repression was observed in both the biofilm and planktonic states. EMSAs were also performed to determine whether Crp directly binds to the promoters of these genes. In contrast to Crp binding to the DNA sequence corresponding to the *pla* promoter in a cAMP-dependent manner (Fig. 5A), Crp did not bind to sequences corresponding to the *ybtA* or *fyuA* promoters (Figs. 5B-C). Thus, the control of Ybt by Crp is likely indirect. Crp did not alter expression of *fur* or the RyhB sRNAs, known regulators of iron acquisition (Datasets 1-2).

It is surprising to find Crp-repressed *ybt* genes while other virulence factors, such as *pla* and *psa,* are Crp-activated. Iron acquisition is critical for *Y. pestis* pathogenesis in multiple infectious routes (50) and would have been predicted to be required in the potential iron-limiting environment of the lung. It is possible that Ybt is necessary early in pneumonia and expressed when glucose is plentiful and *crp* expression is low. As the pneumonia progresses to the pro-inflammatory phase, and host cells die of pyroptosis (51), iron is acquired by other means or is more available and Crp turns off expression of the Ybt genes indirectly.

### Crp is required for growth on non-glucose sugars and directly binds to the *malT* promoter

Another potential important set of genes controlled by Crp during infection are those essential for use of alternative carbon sources as glucose is depleted in the lung. Many of the Crp-activated genes identified by RNA-seq and pathways enriched by GO analysis are involved in metabolism or transport (Datasets 1 and 3). This is not surprising giving the historical role of Crp in regulating the *lac* operon and genes for alternative sugar metabolisms (52). In our RNA-seq data, we found that Crp activated expression of *ptsG,* which is a gene required for acquiring glucose during pneumonia and is also important during bubonic plague (26). Crp was also found to increase expression of *malT,* the transcriptional activator of maltose metabolism (53). Genes *ptsG* and *malT* were confirmed to be controlled by Crp as transcript levels were lower in the *Δcrp* mutant compared to *Y. pestis* by qRT-PCR (Figs. 4E-F), but transcript levels were not increased when grown in glycerol. This regulation was direct as EMSAs demonstrated Crp directly bound to the promoter sequences for *ptsG* and *malT* (Figs. 5D-E). In addition, the binding of Crp to these promoters required cAMP.

The direct regulation of the *malT* promoter by Crp suggested maltose could be an important alternative to glucose to support growth of *Y. pestis*. Indeed, while *Y. pestis* grew well in TMH with maltose (Fig. 6A), the Δ*crp* mutant did not grow with maltose as the sole carbon source. Deletion of *malT* also reduced growth, although this mutant grew better than Δ*crp*, suggesting Crp activates expression of additional genes involved in maltose metabolism. Complementation of *crp* or *malT* restored growth of the Δ*crp* and Δ*malT* mutants, respectively. In addition to maltose, *Y. pestis* grew better in TMH supplemented with galactose compared to TMH alone (Fig. 6B). Genes for galactose catabolism were not significantly different between *Y. pestis* and Δ*crp,* but genes in the *araF* operon for arabinose catabolism are dependent on *crp* for expression (Dataset 1). In TMH supplemented with glucose, *Y. pestis* formed a biofilm at the air-liquid interface that resulted in lowered observed growth in this assay (Fig. S2A). Such robust biofilm formation was not observed in the presence of the other carbon sources. In contrast, Δ*crp* only grew in TMH with glucose (Fig. 6C), suggesting Crp-activated genes are necessary for metabolism of non-glucose carbon sources. However, it is unknown which carbon sources *Y. pestis* uses in the lungs after glucose is consumed.

**FIG. 5.**
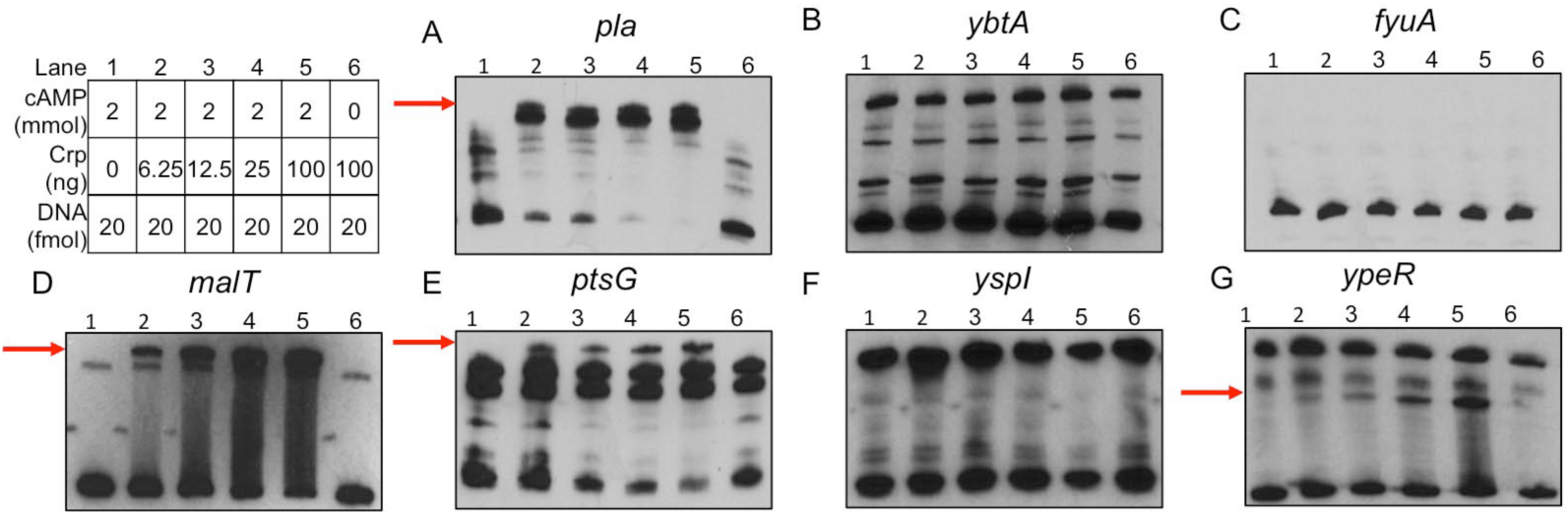
Crp binds to *pla*, *malT*, *ptsG*, and *ypeR* promoter sequences. EMSAs using purified Crp protein incubated with DNA fragments corresponding to promoters for (A) *pla*, (B) *ybtA*, (C) *fyuA*, (D) *malT*, (E) *ptsG*, (F) *yspI* and (G) *ypeR.* Red arrows denote shifted band present in lanes containing Crp protein and cAMP. Table denotes concentrations of Crp protein, cAMP, and promoter DNA used in each reaction.

**FIG. 6.**
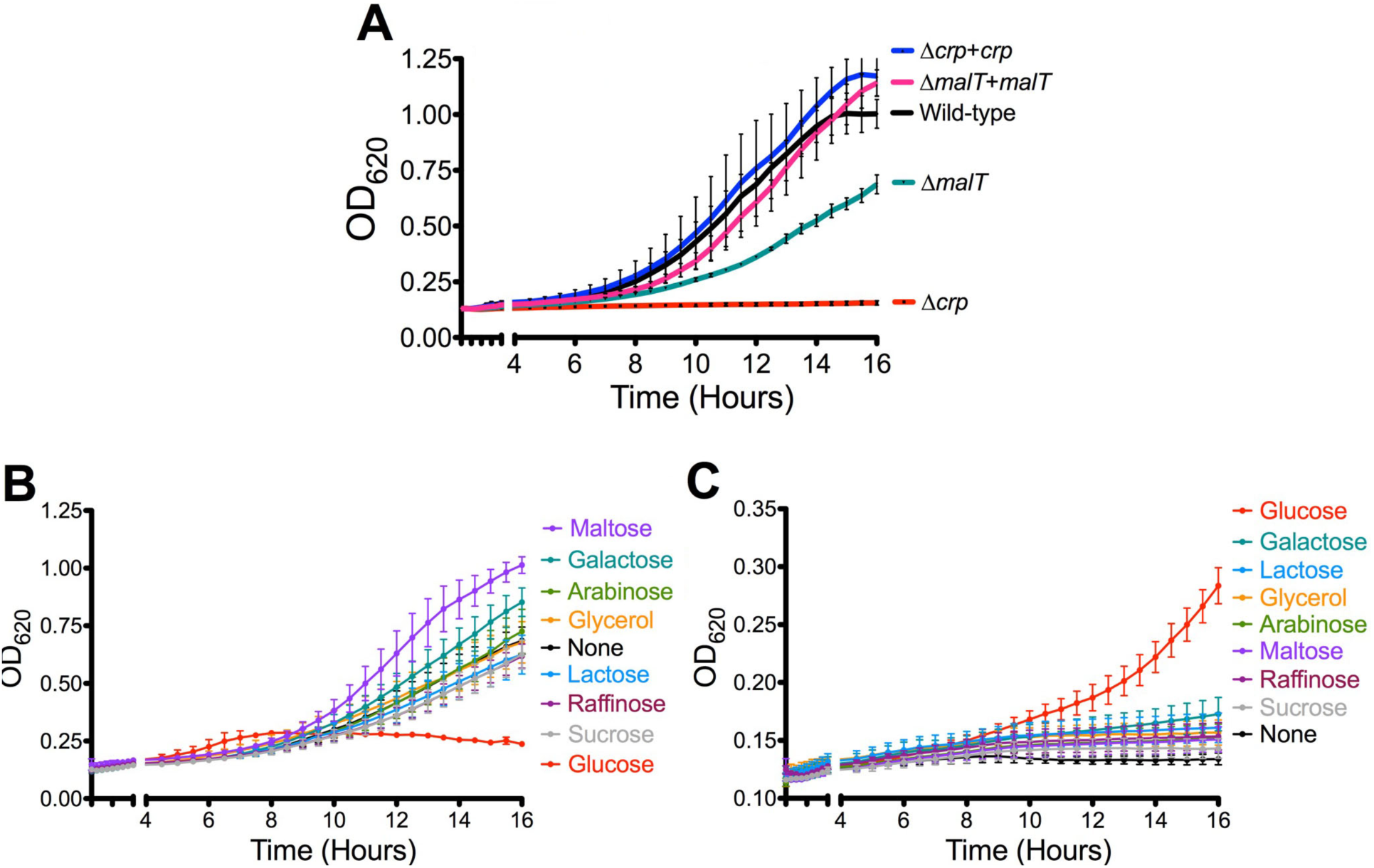
Crp is required for growth on non-glucose sugars. (A) Growth of *Y. pestis* and denoted strains on TMH + 0.2% maltose for 16 hours at 37°C. Growth of *Y. pestis* (B) and *Y. pestis* Δ*crp* (C) in TMH supplemented with no carbon source or 0.2% of the indicated carbon sources for 16 hours at 37°C. Data are combined from three independent experiments and error bars represent the mean and SEM.

### Crp directly and indirectly regulates expression of quorum sensing genes

A final class of potentially important Crp regulated genes identified in this study are those associated with the LuxIR-based acyl-homoserine lactone (AHL) quorum sensing systems. The gene pairs are 100% identical on the *Y. pestis* genome with the AHL synthetase genes (*yspI* and *ypeI*) convergently transcribed with the receptors (*yspR* and *ypeR*) (Fig. 7A). Crp positively controlled expression of both the AHL synthetase genes and the AHL receptors (Table 2). qRT-PCR demonstrated expression of *yspI* and *ypeR* was increased in glycerol and *Y. pestis* compared to Δ*crp,* similar to the positive control *pla* (Figs. 4A,G-H). Moreover, expression of the AHL synthetase, *yspI* was further increased during biofilm growth. A cAMP-Crp dependent shift of the DNA sequence corresponding to the *ypeR* promoter was observed by EMSA (Fig. 5E). However, Crp did not bind to *yspI* promoter sequence suggesting regulation of this gene is indirect (Fig. 5F).

**FIG. 7.**
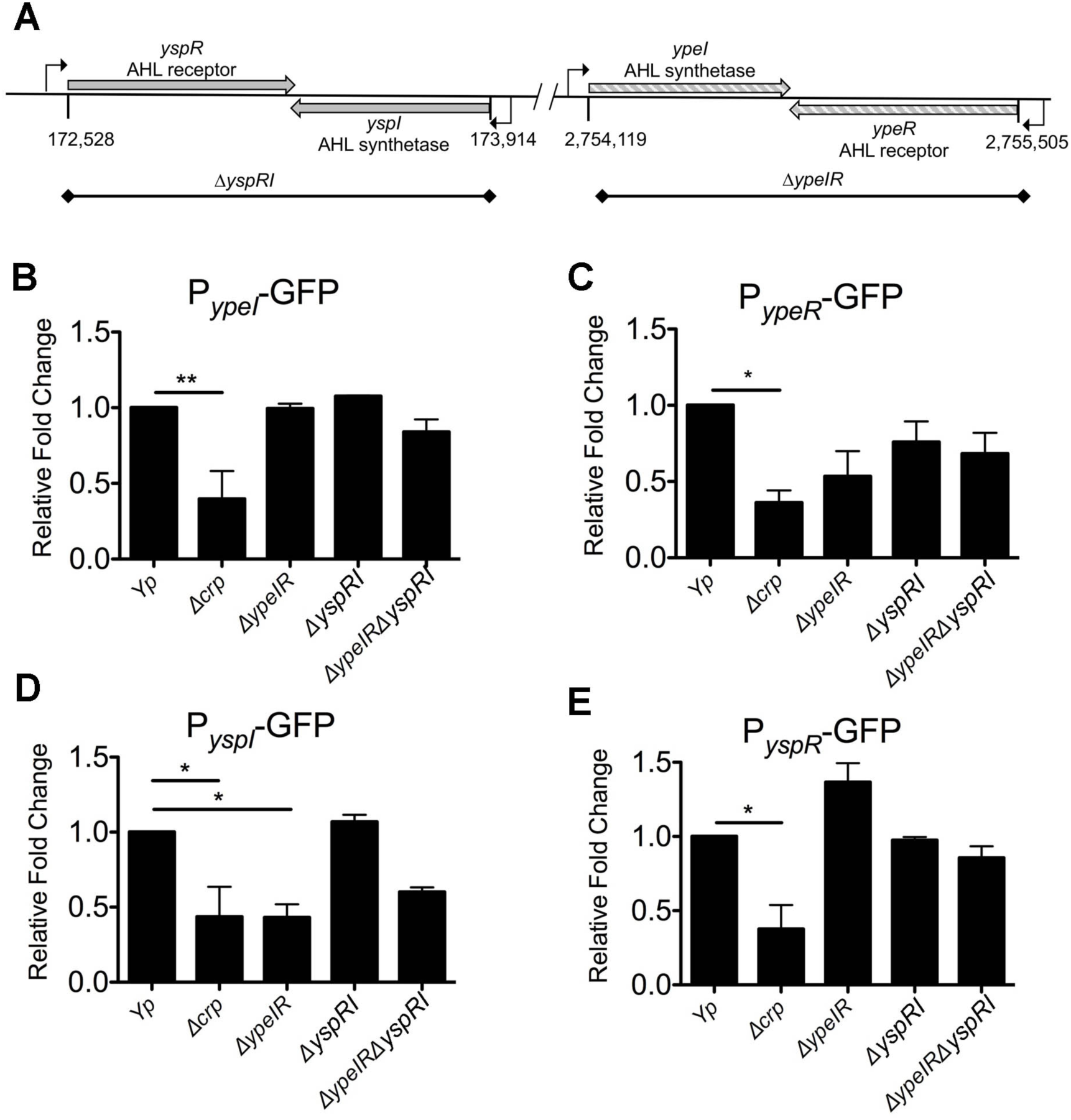
Crp directly and indirectly regulates expression of *Y. pestis* quorum sensing. (A) Schematic of arrangement of *yspIR* and *ypeRI* loci in genome of *Y. pestis* CO92. GFP reporter assays using the (B) P*_ypeI_*-GFP, (C) P*_ypeR_*-GFP, (D) P*_yspI_*-GFP, and (E) P*_yspR_*-GFP reporters integrated into *Y. pestis* and Δ*crp*, Δ*ypeIR, ΔyspIR*, and *ΔypeIRΔyspIR* mutant strains. Background fluorescence was subtracted and remaining signal was normalized to wild-type *Y. pestis* that was set to 1. Data represent the mean and SEM from three independent experiments. \**p*<0.05 and \*\**p*<0.01 from one-way ANOVA with Bonferroni’s multiple comparison test.

In *Y. pseudotuberculosis,* expression of the same quorum sensing genes are auto-regulated, regulated by each other, and Crp-activated (31, 54). Both sets of synthetases and receptors are expressed from separate promoters antiparallel to each other (Fig. 7A). To determine if a similar pattern occurs in *Y. pestis,* GFP-reporters containing the promoter regions of *yspR*, *yspI*, *ypeI*, and *ypeR* were integrated into *Y. pestis,* Δ*crp,* or *Y. pestis* with deletion of *ypeI* plus *ypeR (*Δ*ypeIR), yspR* plus *yspI (*Δ*yspRI),* or *Y. pestis* with deletion all four AHL genes (Δ*yspRI*Δ*ypeIR*). Expression of all four reporters was reduced approximately 50% in Δ*crp,* confirming RNA-seq and qRT-PCR data that Crp stimulates expression of the *ype* and *ysp* AHL genes (Figs. 7B-E). Expression of the P*_ypeI_-*GFP reporter showed no dependence upon *ysp* or *ype* genes. The P*_ypeR_-*GFP reporter expression was reduced in Δ*ypeIR* and Δ*yspRI*Δ*ypeIR* suggesting *ypeR* expression may be positively auto-regulated, but the differences were not statistically significant. The exact opposite occurs in *Y. pseudotuberculosis* wherein the *ypeIR* (*ypsIR*) genes repress their own expression at 37°C (54). Expression of the P*_yspI_-*GFP reporter was reduced two-fold in *Y. pestis* Δ*ypeIR* suggesting the reduction in *yspI* expression in *Y. pestis* Δ*crp* could be in part due to reduced expression of *ypeR* (Fig. 4G). Similarly, the *yspI* homolog in *Y. pseudotuberculosis* is dependent on *ypeR* (*ypsR* in *Y. pseudotuberculosis*) (54) and expression of *ypeIR* and *yspIR* homologs in *Y. pseudotuberculosis* YPIII may also be Crp-regulated (31).

All total, these data suggest direct regulation by Crp at the *ypeR* promoter and indirect regulation of *yspI* through *ypeR* resulting in stimulation of genes in glucose limited media in biofilm. Despite the coding sequence of these genes being 100% identical to each other, the quorum sensing genes are regulated differently between and within *Y. pestis* and *Y. pseudotuberculosis*.

### Crp regulates production of AHLs *in vitro*

Given the decrease in expression of *yspI* and *ypeI* in Δ*crp,* production of AHLs should also be reduced. Cell-free supernatants from *Y. pestis* strains were collected during planktonic growth *in vitro* and were used to stimulate the *Rhizobium radiobacter* AHL bioreporter. AHLs produced by *Y. pestis* increased with cell density during exponential phase as expected for quorum sensing (Fig. S3A). In contrast, AHL production in Δ*crp* increased only slightly despite a six-fold increase in bacterial density. Because growth rate of Δ*crp* is reduced, AHL concentrations were also compared at similar cell densities between strains. Even when accounting for cell density, AHL production was reduced in the *Δcrp* strain (Fig. 8A). These data suggest the reduction in AHL production is due to reduced *ypeI* and *yspI* expression and not differences in cell density or growth of Δ*crp*. To determine if Crp alters production of specific AHLs, extracts were developed by thin-layer chromatography (TLC). Three spots were observed in all three strains corresponding to 3-oxo-C6-HSL, C6-HSL, and C8-HSL (top to bottom) as reported previously for *Y. pestis* (55). However, the areas of each of the spots from Δ*crp* were smaller (Fig. 8C). No spots were detectable in extracts collected from Δ*ypeIR*Δ*yspRI* (ΔAHL, Fig. 8D).

**FIG. 8.**
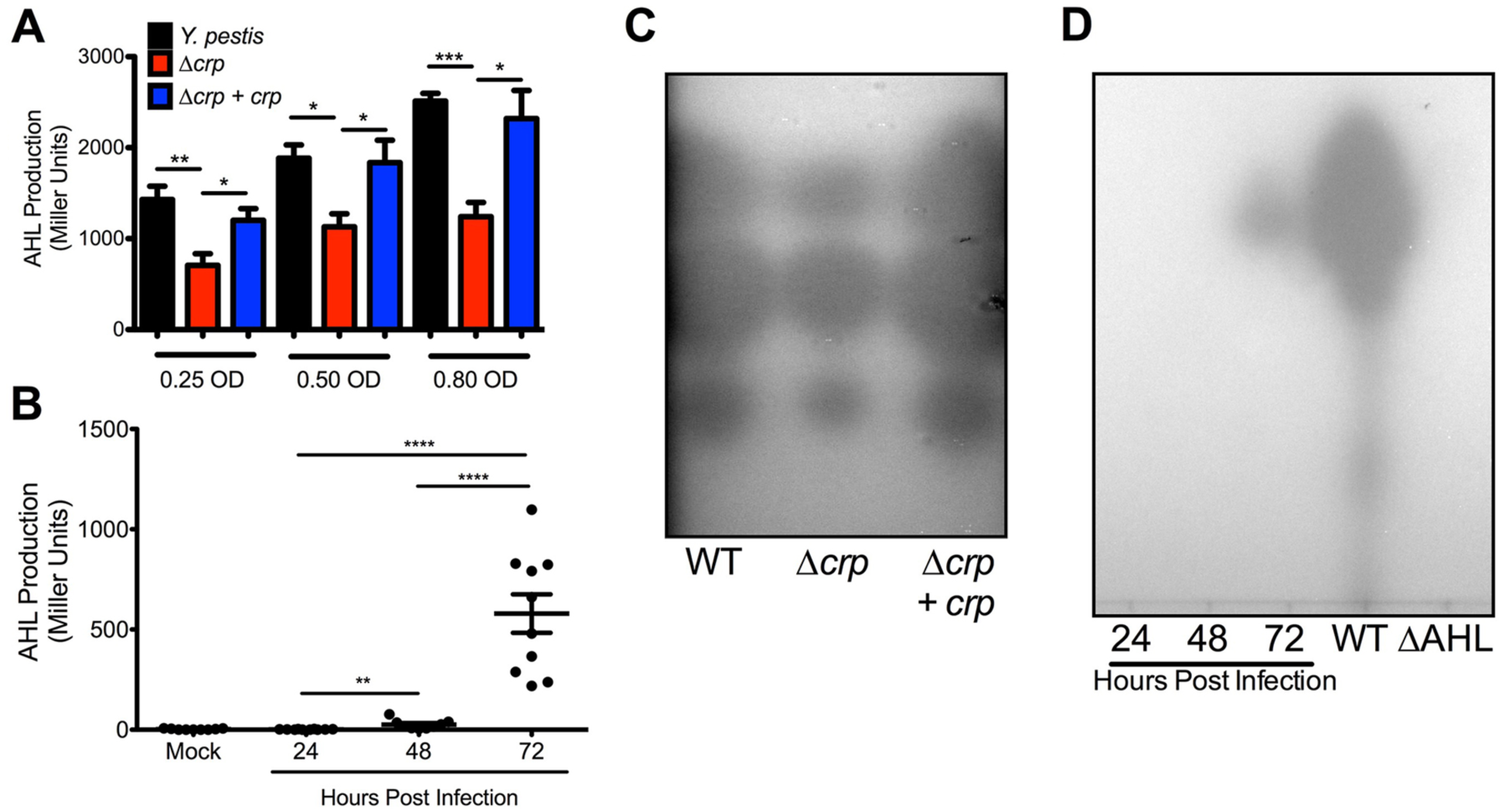
AHL production requires Crp and increases during pneumonic plague. (A) Comparison of AHL production between *Y. pestis, Δcrp*, and *Δcrp+crp* strains at equivalent cell densities. (B) Concentrations of AHLs in BALF collected from mice infected with *Y. pestis* intranasally. (C) Representative TLC plate of AHL extracts collected from *Y. pestis, Δcrp*, and Δ*crp+crp* strains grown to 0.80 OD_620_ (*Y. pestis* and Δ*crp+crp* 6 hours, *Δcrp* after 12 hours). From top to bottom, intensity of the AHL spots from Δ*crp* were 77%, 83%, and 76% of the area of wild-type corresponding to *N*-(3-oxohexanoyl)-L-homoserine, *N*-hexanoyl-L-homoserine, and *N*-octanoyl-L-homoserine lactones respectively (55). Data are representative of three-independent experiments. Spot intensities were calculated in FIJI. (D) Representative TLC plate of AHL extracts collected from BALF samples. \**p*<0.05, \*\**p*<0.01, \*\*\**p*<0.001, and \*\*\*\**p*<0.0001 from Student’s *t*-test.

To determine whether AHL molecules are produced during lung infection, bronchoalveolar lavage fluid (BALF) was collected from mice infected with fully virulent *Y. pestis.* AHLs were undetectable in BALF from mock-infected mice and mice at 24 hour post infection (hpi) (Fig. 8B). Only low levels of AHLs were detectable in BALF 48 hpi, while BALF 72 hpi contained a 10-fold increase in AHLs. Similar trends were present for AHLs from BALF developed by TLC plates (Fig. 8D). The significant increase in AHLs at 72 hpi correlates with the timeframe during pneumonia when *Y. pestis* depletes glucose from the lungs, forms large biofilms, and as a result, expresses *crp* to a high level (Fig. 1A)(26). The formation of biofilms in the lungs may provide a favorable environment for amplification of quorum sensing-based gene regulation. Indeed, expression of *yspI in vitro* was more highly expressed in glycerol-grown biofilms compared to glucose-grown biofilms (Figs. 4H-I). Taken together these data provide strong evidence for Crp-dependent production of AHLs and induction of quorum sensing during lung infection. Transcriptome studies have linked *Y. pestis* quorum sensing mutants to defects in multiple behaviors. Deletion of both synthetase-receptor pairs impairs the ability for *Y. pestis* to make biofilms and metabolize maltose (55, 56). Both of these phenotypes are also Crp-dependent (Fig. 6)(12), and suggest that connecting the Crp regulon to the quorum sensing regulon may allow *Y. pestis* to cooperatively regulate gene expression for when glucose is gone and the bacterial density is high.

While multiple measurements indicated Crp regulates AHL-based quorum sensing, autoinducer-2 (AI-2) or LuxS-based quorum sensing is not Crp-regulated in the planktonic state (Dataset 1 and Table 2). AI-2 secretion was not significantly different between *Y. pestis* and *Δcrp* at equivalent cell densities in planktonic culture (Fig S3B-C). In contrast to AHLs, the concentration of AI-2 in BALF samples increased in a step-wise manner from 24 to 48 to 72 hpi as *Y. pestis* density grows within the lungs even prior to increased expression of *crp* (Fig. S3D) (26). These data suggest AI-2 production is not dependent on Crp, but rather bacterial cell density increases between 24 and 72 hpi.

## Conclusions

In this study, we solved the crystal structure of *Y. pestis* Crp while determining its regulon under planktonic and biofilm conditions. While structurally similar to *E. coli* Crp, the regulation of the *crp* gene differs amongst pathogenic *Yersinia* spp., which colonize distinct environmental niches. Overall, transcription profiling conducted in this study revealed comprehensive insight into how *Y. pestis* adapts to growth in the biofilm state and the role of Crp and carbon sources during this growth. This is specifically relevant in the lungs, when Crp, through sensing of depletion of glucose, may serve as a switch between turning on or off multiple behaviors of *Y. pestis* as disease in the lung progresses (26)(Fig. 9).

**FIG. 9.**
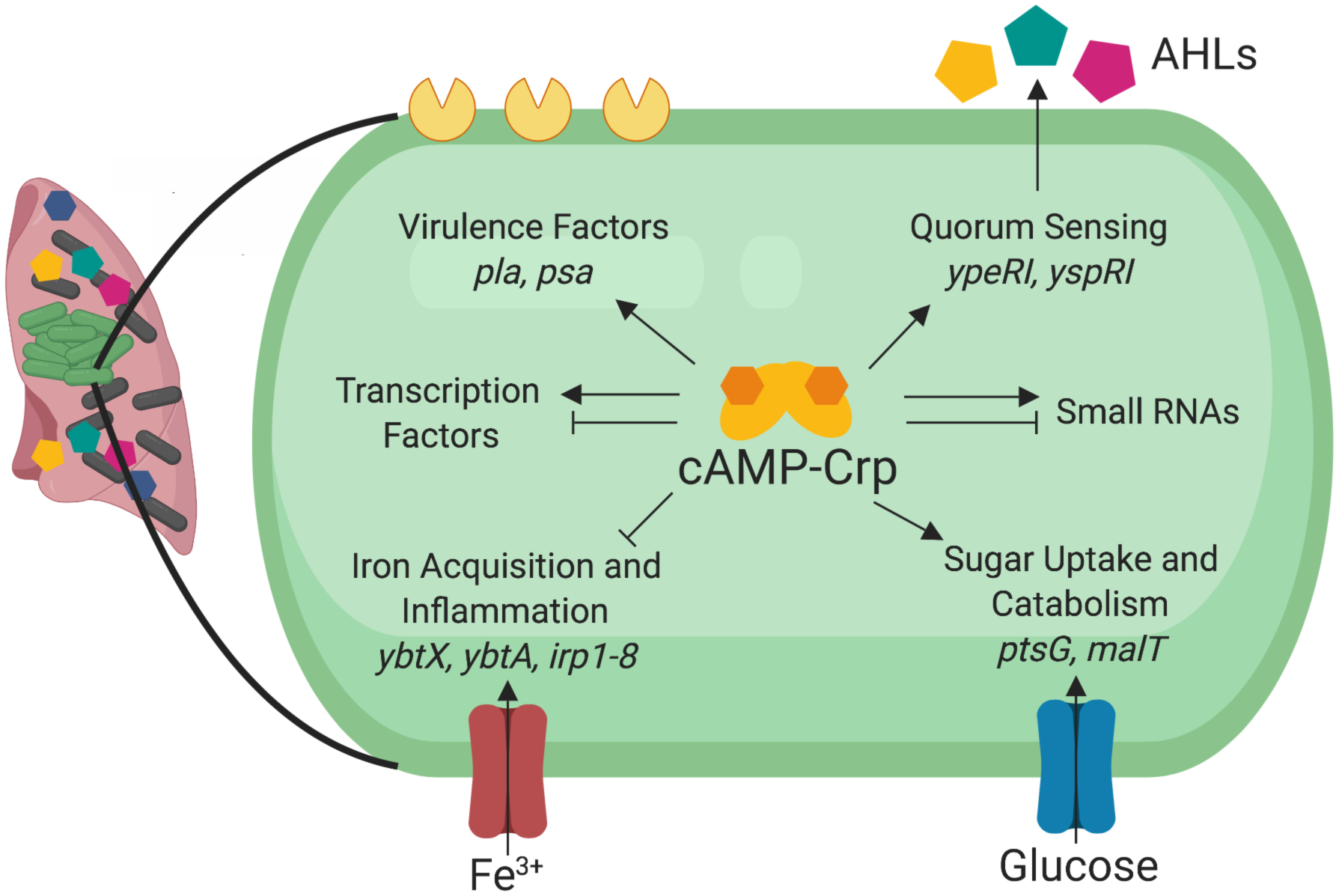
Model of identified Crp-activated genes and pathways in *Y. pestis.* Crp requires cAMP in order to regulate multiple behaviors of *Y. pestis* including virulence, quorum sensing, iron acquisition and inflammation, and sugar uptake and catabolism. cAMP-Crp also regulates expression of hundreds of other mRNAs, including transcription factors, and small non-coding RNAs.

A unique result of our transcriptional profiling was the observed indirect repression of Ybt genes by Crp (Table 1 and Table S2). The Ybt and *irp* genes are critical for iron acquisition, virulence, and inflammation during pneumonic plague (48–50). Expression of these genes may be higher up to 48 hpi, when expression and activity of Crp is low (Fig. 1A). After 48 hpi, Crp activates a second set of virulence factors such as *pla* and *psa* as disease progresses correlating with the biphasic nature of pneumonic plague (23, 27, 32). The regulation of iron-acquisition genes is unique to *Y. pestis* compared to *Y. pseudotuberculosis.* Even though these closely related species share a 100% identical Crp, the expression of the *crp* gene and environments inhabited by these species may afford different sets of regulated genes.

We expand the pool of directly Crp-activated genes in *Y. pestis* to include genes for quorum sensing and alternative metabolisms. As glucose becomes a limiting resource, genes for carbohydrate catabolism and uptake are activated to support growth of *Y. pestis.* Within biofilms of the lungs, AHL-dependent quorum sensing is likely amplified due to the close proximity of cells. Here, we report production of AHLs in the lungs of plague-infected mice correlating with the time expression of *crp* is highest and *Y. pestis* is growing within biofilms of the lungs. Thus, not only is Crp directly regulating its own genes, but other transcriptional regulators and regulons (Fig. 9). These additional pathways may direct additional *Yersinia* behaviors and may serve as a link to the indirect regulations observed in our dataset. Future experimentation on differentially expressed mRNAs and sRNAs identified in this study will provide additional information on biofilm-regulated and Crp-regulated genes and behaviors in *Y. pestis* and its importance during pneumonia.

## MATERIALS AND METHODS

### Bacterial strains, media, and growth conditions

Bacterial strains, plasmids and primers and are listed in Table S4. *Y. pestis* was passaged on Difco^TM^ brain heart infusion (BHI) agar (Becton Dickinson) and into 2 mL of BHI broth or the defined media, TMH (57), supplemented with 0.2% glucose for overnight growth at 26°C. Unless otherwise stated overnight cultures in TMH were subcultured at an optical density 620 nm (OD_620_) = 0.1 into 10 mL of TMH with 0.2% glucose in 125-mL Erlenmeyer flasks incubated at 37°C with shaking at 250 rpm for measuring planktonic and biofilm cell differences. Planktonic cells were removed from the flask with a serological pipette. Biofilms were scraped and resuspended in phosphate-buffered saline (PBS) warmed to 37°C. For growth assays, overnight cultures were subcultured at an OD_620_ = 0.1 into 200 µL of fresh TMH with 0.2% of indicated carbon sources at 37°C in a 96-well plate. Absorbance was measured with a Molecular Devices Spectramax M5 microplate reader. *E. coli* strains were passaged on Luria-Bertani (LB) agar or in LB broth at 37°C. *Vibrio harveyi* MM32 was cultured in ATCC medium 2034 or LB with kanamycin. *R. radiobacter* was passaged on BHI with gentamicin. Ampicillin (100µg/mL), kanamycin (50µg/mL), and gentamicin (15µg/mL) were added as necessary.

### Construction of new strains and plasmids

*Y. pestis* containing deletions in *crp, ypeIR, yspRI,* and *luxS* were generated by lambda red recombination (6). The *ΔypeIR*Δ*yspRI* double mutant was generated by deletion of the *ypeIR* locus from *ΔyspRI*. *Y. pestis* Δ*crp* and Δ*malT* were complemented *in trans* by cloning the *crp* or *malT* codons plus 500 base pairs (bp) flanking the genes into pUC18-R6k-miniTn7t by Gibson assembly. The resulting plasmid was integrated into the Tn*7* site (58). GFP reporters were also integrated into the Tn*7* site. 500 bp upstream of the promoters for *yspI, yspR, ypeI,* and *ypeR* were amplified by PCR and adjoined to the codons of *gfp* by overlap extension PCR as described previously (35).

### RNA extraction, sequencing, and qRT-PCR

Total RNA was isolated from *Y. pestis* cultures as described previously (59) except that RNA protect reagent (Qiagen) was added to planktonic or biofilm cells in PBS. RNA was treated with 1 µL of Turbo DNAse (Invitrogen Life Technologies) for 30 minutes at 37°C followed by addition of another 1 µL for 30 minutes following manufacturer’s instructions.

Next-generation RNA-sequencing was carried out at the Northwestern University NUSeq Core Facility. Total RNA was checked for quality and quantity on Agilent Bioanalyzer 2100 and Qubit fluorometer. The Illumina TruSeq Stranded Total RNA Library Preparation Kit was used to prepare sequencing libraries from 500 ng of total RNA samples according to manufacturer’s instructions without modifications. This procedure includes rRNA depletion with Ribo-Zero rRNA Removal Kit (Bacteria), remaining RNA purification and fragmentation, cDNA synthesis, 3’-end adenylation, Illumina adapter ligation, library PCR amplification and validation. Illumina NextSeq 500 Sequencer was used to sequence the libraries with the production of single-end, 75 bp reads.

The quality of DNA reads, in fastq format, was evaluated using FastQC. Adapters were trimmed, and reads of poor quality or aligning to rRNA sequences were filtered. The cleaned reads were aligned to the *Y. pestis* genome (NC_003143.1) and plasmids pPCP1 (NC_003132.1) and pMT1 (NC_003134.1) using STAR (60). Read counts for each gene were calculated using htseq-count (61) in conjunction with a gene annotation file for *Y. pestis*. Read counts for sRNA were obtained using bedtools using the annotated locations from Schiano et al. (33). Normalization and differential expression were determined using DESeq2 (62). The cutoff for determining significantly differentially expressed genes was an FDR-adjusted *p*-value less than 0.05. Raw and processed data were deposited into the GEO database at the NCBI (accession number: GSE135228).

For validation of RNA-sequencing results, qRT-PCR was performed with the SuperScript III Platinum Syber Green One-Step qRT-PCR Kit (ThermoFisher Scientific) using 25 ng of DNased RNA following the manufacturer’s instructions on a BioRad iQ5 cycler with melt curve analysis.

### Purification of *Y. pestis* Crp

The amplified *crp* gene corresponding to residues 1-210 from *Y. pestis* CO92 (www.csgid.org IDP97063) was cloned by Gibson assembly into the SspI site of expression vector pMCSG7 (63), which contains an N-terminal polyhistidine tag followed by the tobacco etch mosaic virus (TEV) protease cleavage site and the start codon of the *crp* gene. The resulting plasmid was transformed into *E. coli* BL21(DE3)(Magic). The bacteria were grown at 37°C, 200 r.p.m. in 3 L of Terrific Broth until OD_600_ nm reached 1.6. Protein expression was induced with 0.6 mM IPTG and cells were grown overnight with shaking reduced to 180 r.p.m. and temperature to 22°C. Cells were harvested by centrifugation. The resulting cell pellet was re-suspended in 120 ml of lysis buffer [10 mM Tris-HCl (pH 8.3), 500 mM NaCl, 1mM Tris (2-carboxyethyl) phosphine (TCEP), 10% (v/v) glycerol, 0.01% (v/v) IGEPAL CA630, EDTA-free protease inhibitors (64) 1 tablet/ 100 ml buffer] and the suspension was frozen at -20°C until purification. The frozen suspension was thawed under the cold running water, sonicated and centrifuged. The protein was purified in two steps using nickel (II) affinity chromatography followed by size exclusion chromatography as described (65). The poly-histidine tag was removed by incubation of the tagged protein with the recombinant TEV protease for overnight at 20°C. The resulted 58 mg of pure protein was at a final concentration of 13.7 mg/ml.

### Structure determination of *Y. pestis* Crp

For crystallization screening, we used protein solution with concentration of 6.6 mg/ml in 10 mM Tris-HCl (pH 8.3), 500 mM NaCl, 5 mM TCEP, and 1 mM cAMP. Crystallization drops in 1:1 ratio (protein: reservoir solution) were equilibrated against 96 conditions/screen using commercially available PACT, PEG’s and PEG’s II Suites (Qiagen). Diffraction quality crystals of Crp grew from PACT Suite condition H12.

Prior to flash-cooling in liquid nitrogen, crystals of Crp were transferred into a 5 µl drop of reservoir solution they grew from. Data were collected on the LS-CAT 21-ID-F beamline at the Advanced Photon Source (APS) at Argonne National Laboratory. A total of 350 images were indexed, scaled and integrated using HKL-3000 (66). Data collection and data processing statistics are listed in Table S1. The structure of *Y. pestis* Crp in complex with cAMP was solved by Molecular Replacement using Phaser (67) from the CCP4 suite (68). The structure *E. coli* Crp (PDB entry 3RYP) was used as a search model. The initial solution went through several rounds of refinement in REFMAC v.5.5, residues were mutated and cAMP was added in Coot (69). Water molecules were generated using ARP/wARP (70) and the model refinement was continued in REFMAC. Translation–Libration–Screw (TLS) groups were created by the *TLSMD* server ((71); http://skuldbmsc.washington.edu/~tlsmd/) and TLS corrections were applied during the final stages of refinement. *MolProbity* (Chen *etal.*,2010; http://molprobity.biochem.duke.edu/) was used for monitoring the quality of the model during refinement and for the final validation of the structure. The final model and diffraction data were deposited to the Protein Data Bank (https://www.rcsb.org/) with the assigned PDB entry 6DT4. The final model consists of two polypeptide chains, which form a dimer. Chain A contains Crp residues 6-208 and chain B contains residues 9-208. There is one cAMP molecule bound to each monomer, 3 chloride ions, and 292 water molecules in the crystal structure. Refinement statistics and the quality of the final model are summarized in Table S1. Molecular graphics and alignment with *E. coli* Crp bound to cAMP (PDB ID 4R8H (30)), were performed with the UCSF Chimera package.

### Gel shift assays

EMSAs were carried out using the LightShift^TM^ Chemiluminescent EMSA Kit (ThermoFisher Scientific). Biotinylated primers were used to PCR-amplify gene-specific promoters (Table S5). PCR products were concentrated and gel purified using Wizard SV Gel and PCR cleanup protocol (Promega). Binding reactions consisted of the manufacturer’s binding buffer, 100 ng Poly (dI:dC), and 20 fmol of biotinylated promoter fragment in 20 µL. Reactions contained increasing amounts of purified Crp protein (6.25-100 ng), and cAMP (2 mmol, Sigma). Binding reactions were carried out following manufacturer’s instructions. Nylon membranes (Hybond-N+, 30 cm) were developed using X-ray film.

### GFP reporter assays

For GFP reporter assays, 2 mL overnight *Y. pestis* cultures in TMH with 0.2% glucose were subcultured at an OD_620_ = 0.1 into 2 mL of TMH incubated at 37°C in a rotary drum for six hours. Cultures were diluted to OD_620_ of 0.25-0.40 and 200 µL aliquots were added in duplicate to a 96-well plate. Fluorescence was read on a Tecan Safire II plate reader and reported as previously described (6).

### AHL and AI-2 bioreporter assays

*R. radiobacter* carrying pZLR4 was grown overnight at 30°C. Cultures were diluted to OD_620_ = 0.1 in 1 mL of BHI. 1 µL of filter-sterilized supernatants of *Y. pestis* or 10 µL of filter-sterilized BALF, previously collected from mice intranasally infected with wild-type *Y. pestis* (26), was added. Tubes were incubated at 30°C with shaking at 250 RPM for 5 hours. Cultures were pelleted and resuspended in 500 µL of Z-buffer (60 mM Na_2_HPO_4_, 40 mM NaH_2_PO_4_, 10 mM KCl, 1 mM MgSO_4_, pH = 7.0). 25 µL of chloroform and 12.5 µL of 0.1% SDS was used to lyse the bacteria for 5 minutes. 100 µL of 2-nitrophenyl-β-D-galactopyranoside (ONPG, 4 mg/mL) was added to initiate the galactosidase reaction, which was stopped by the addition of 250 µL of 1 M Na_2_CO_3_. The reaction mixtures were centrifuged for 5 minutes at 13,000 RPM and 500 µL was added to a cuvette to record the OD420. Relative beta-galactosidase activity was expressed as the ratio of OD_420_/OD_620_.

For TLC, 100 µL of culture supernatants or BALF were extracted twice with 100 µL of ethyl acetate and concentrated to 20 µL in a 60 Hz Savant SpeedVac DNA 100 concentrator (ThermoFisher Scientific). 1 µL of culture supernatants or 7.5 µL of BALF samples were spotted onto aluminum-backed C18-W silica plates (Sorbent Technologies) and developed in 60% methanol 40% water as described previously (55).

To measure AI-2 concentrations, overnight cultures of MM32 were diluted 1:500 and grown at 30°C for 1 hour. 180 µL was added to a 96-well plate with either 20 µL of culture supernatant or BALF. Plates were incubated at 30°C with shaking for 5 hours. Optical density and luminescence were recorded on a Molecular Devices Spectramax M5 microplate reader. Reported data are from 3 hours incubation (culture supernatants) and 4 hour incubation (BALF).

### Measurement of intracellular cAMP concentration

Overnight cultures of *Y. pestis* were diluted to OD_620_ = 0.1 in TMH with 0.2% glucose or 0.2% glycerol and grown for six hours at 26°C or 37°C. Alternatively *Y. pestis* was grown in TMH overnight in 125-mL Erlenmeyer flasks to form biofilms. 0.2 OD_620_ equivalents were centrifuged and lysed in 0.1M HCl and 0.5% Triton X-100 for 10 minutes. Cellular debris was pelleted by centrifugation and the supernatant was stored at -20°C. The Direct cAMP ELISA kit (72) was used following the acetylation protocol. cAMP concentrations were normalized to protein content measured by the Pierce BCA Protein Assay Kit (ThermoFisher Scientific).

### Statistical Analysis and Graphing

Statistical analyses including Student’s *t* tests and one-way analysis of variance ANOVA with Bonferroni multiple-comparison tests were performed in GraphPad Prism, version 5.0, using a *p* value of <0.05 as a cutoff for significance. Venn diagrams and volcano plots were generated in R (Version 9.0) using the VennDiagram (Version 1.6.20) and tidyverse (Version 1.1.1) packages.

### Gene Ontology Analysis

RNA-seq was analyzed at GeneOntology.org (43, 44) using the GO Enrichment Analysis (77). The PANTHER Overrepresentation Test (Released 2019711) was conducted with the GO Ontology database released 2019-07-03. The reference list used was *Yersinia pestis*.

## SUPPLEMENTAL MATERIAL

**Fig. S1.** Principle-component analysis of RNA-seq dataset. Principle component analysis between the six different conditions and three different replicates.

**Fig. S2.** Biofilm formation after growth in TMH. Crystal violet staining of (A) *Y. pestis* and (B) *Y. pestis Δcrp* biofilms formed after 16 hours of growth in TMH supplemented with indicated carbon sources. Wells were stained as described previously (35).

**Fig. S3.** Crp does not regulate AI-2 based quorum sensing.

**Table S1.** X-Ray Data Collection and Refinement Statistics of *Y. pestis* Crp

**Table S2.** Crp-activated and repressed genes.

**Table S3.** List of bacterial strains and plasmids used in this study.

**Table S4.** List of oligonucleotides used in this study.

**Dataset S1.** List of differentially expressed mRNAs in *Y. pestis* CO92 between all test conditions.

**Dataset S2.** List of differentially expressed sRNAs previously identified in *Y. pestis* CO92 between all test conditions.

Dataset S3. Gene Ontology

## ACKNOWLEDGEMENTS

We would like to thank Dr. Ludmilla Shuvalova and Olga Kiryukhina for expression, purification, crystallization and data collection of the Crp protein. Use of the Tecan Safire II plate reader and access to structure determination software and LS-CAT were provided by the Northwestern University Structural Biology Facility supported by NCICSG P30-CA60553. Chimera is developed by the Resource for Biocomputing, Visualization, and Informatics at the University of California, San Francisco (supported by NIGMS P41-GM103311). This project has been funded in whole or in part with Federal funds from the National Institute of Allergy and Infectious Diseases, National Institutes of Health, Department of Health and Human Services, under contract number HHSN272201700060C and grant number R21AI111018 (to K.J.S.)

