## Supplemental Data for "The Cyclic AMP Receptor Protein Regulates Quorum Sensing and Global Gene Expression in *Yersinia pestis* During Planktonic Growth and Growth in Biofilms"

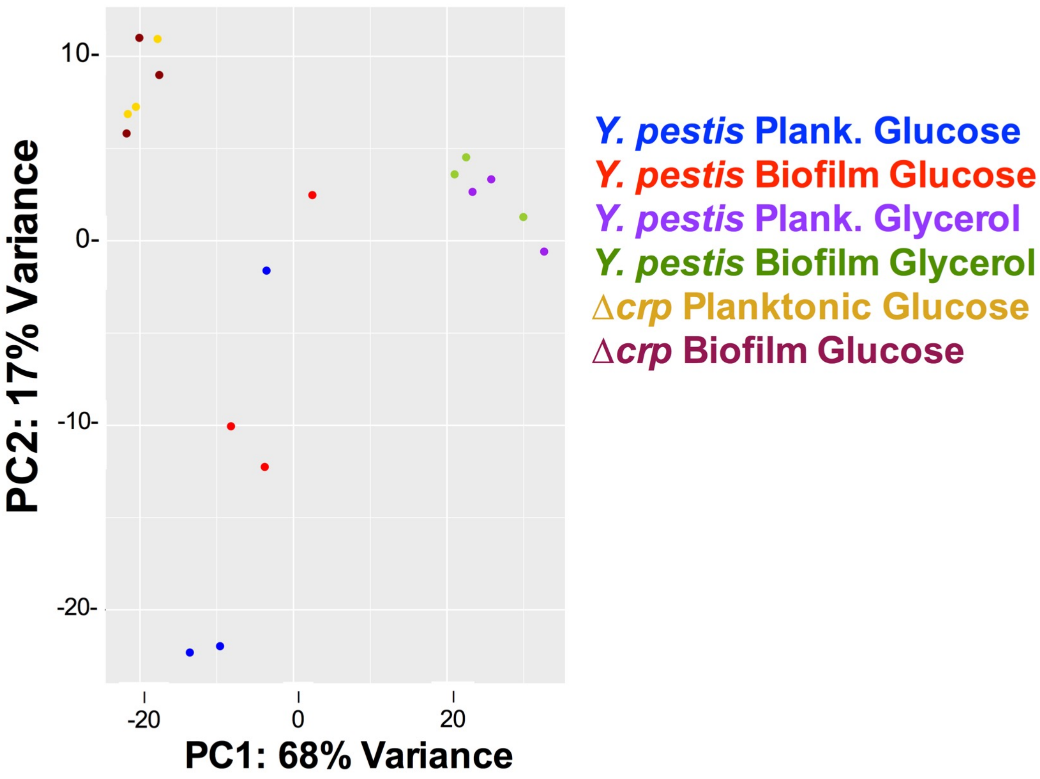


**FIG S1.** Principle-component analysis of RNA-seq dataset. Principle component analysis between the six different conditions and three different replicates.

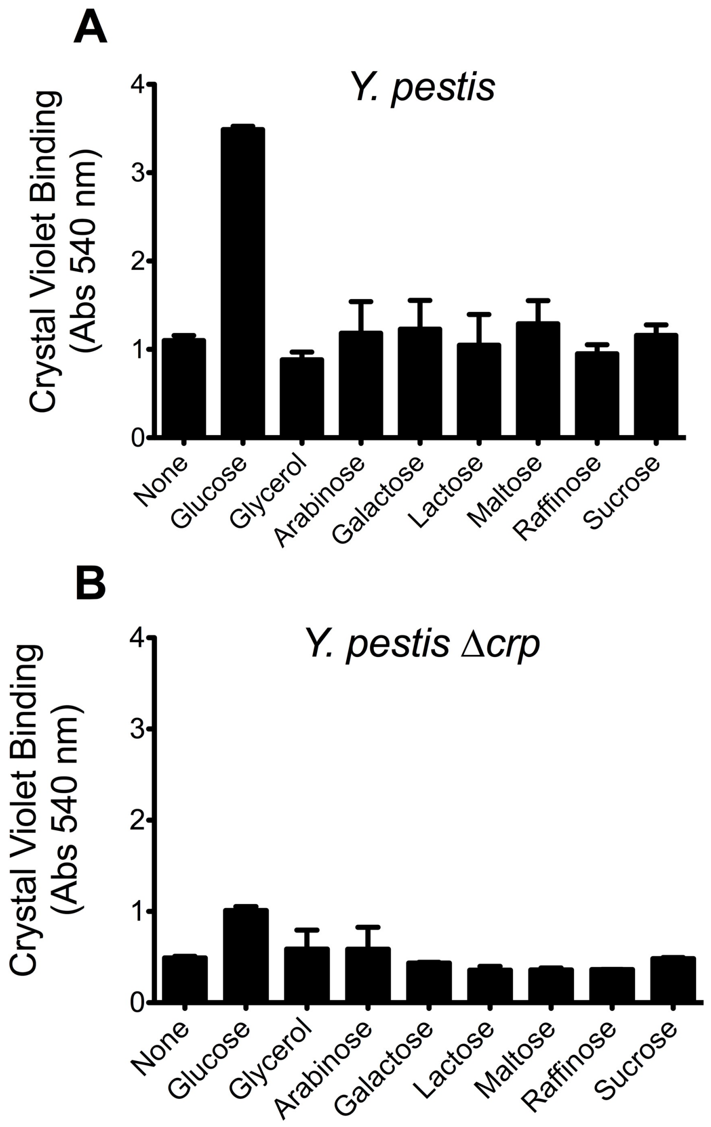


**FIG S2.** Biofilm formation after growth in TMH. Crystal violet staining of (A) *Y. pestis* and (B) *Y. pestis* Δ*crp* biofilms formed after 16 hours of growth in TMH supplemented with indicated carbon sources. Wells were stained as described previously (35).

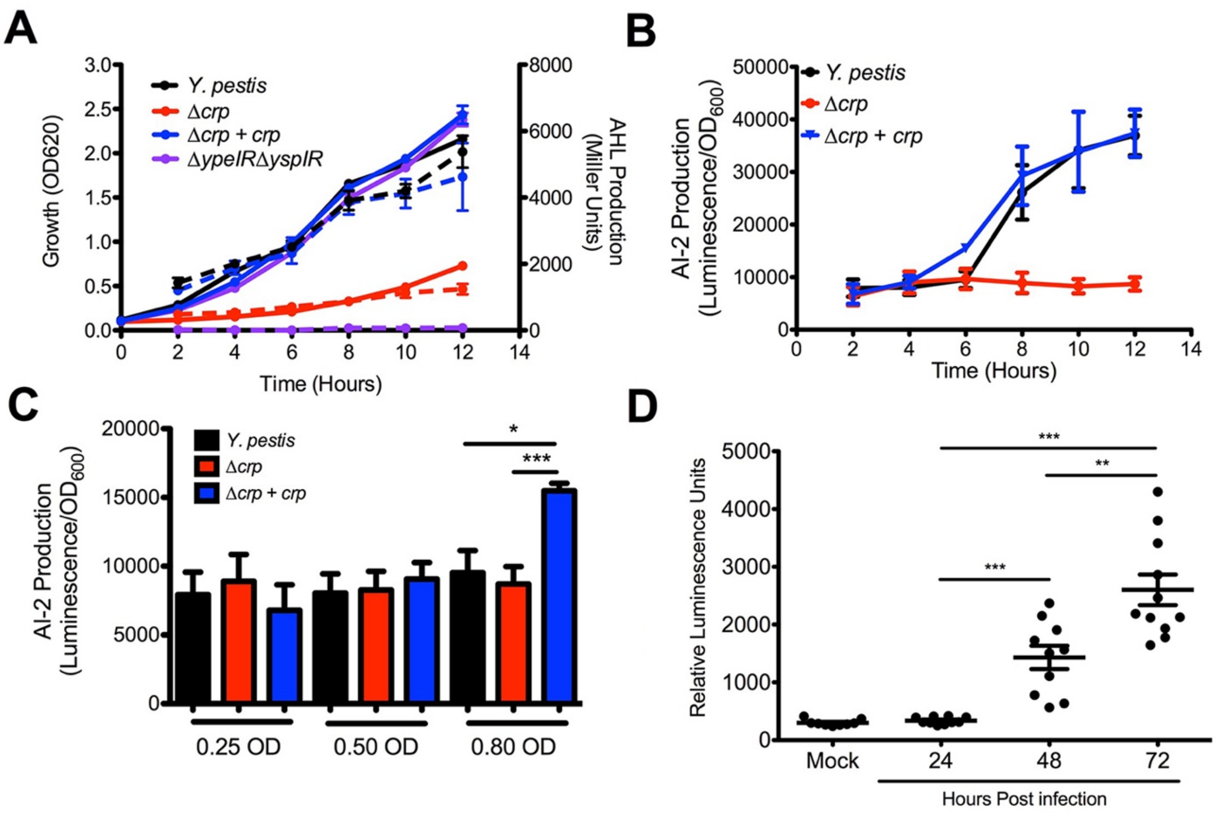


**Fig. S3.** Crp does not regulate AI-2 based quorum sensing. (A) The growth ($OD_{620}$, left *y*-axis, solid lines) and AHL-production (Miller units, right *y*-axis, dotted lines) of *Y. pestis* (black), Δ*crp* (red), Δ*crp*+*crp* (blue), and Δ*ypeIR*Δ*yspIR* (purple) strains. Strains were grown in TMH with 0.2% glucose at 37°C. Every two hours the $OD_{620}$ of cultures were measured and culture supernatants were collected and used to stimulate the *R. rhizobium* bioreporter. (B) AI-2 production measured by the *V. harveyi* MM32 bioreporter from supernatants collected in (A). (C) Comparison of AI-2 production by *Y. pestis,* Δ*crp,* and Δ*crp*+*crp* strains at equivalent cell densities. Data represent the mean and SEM from four independent experiments. (D) Concentrations of AI-2 and in BALF collected from mice infected with *Y. pestis*. Data are combined from two independent animal infections with n=5 at each time point. **p*<0.05, ***p*<0.01, ****p*<0.001, and *****p*<0.0001 from Student's *t*-test.

**TABLE S1** X-Ray Data Collection and Refinement Statistics of *Y. pestis* Crp

| Crystal | *Y. pestis* Crp (PDB ID 6DT4) |
|---|---|
| **Data Collection** | |
| Diffraction Source | Beamline 21ID-F, APS |
| X-ray Wavelength (Å) | 0.97872 |
| Space group) | P 21 21 21 |
| a, b, c (Å) | 53.876, 82.800, 106.350 |
| α, β, γ (°) | 90.00, 90.00, 90.00 |
| Resolution range (Å) | 30.00 – 1.80 (1.83 – 1.80) |
| No. of unique reflections | 44,393 (2,134) |
| Data Completeness (%) | 99.1 (97.3) |
| Multiplicity | 8.5 (7.5) |
| ⟨I/σ(I)⟩ | 30.5 (2.9) |
| $R_{sym}$ (%) † | 5.8 (72.2) |
| Wilson plot B factor (Å$^2$) | 29.9 |
| **Structure Refinement** | |
| Resolution range (Å) | 29.61-1.80 (1.85-1.80) |
| Completeness (%) | 99.0 (96.6) |
| No. of observed reflections | 44349 (3122) |
| No. of $R_{free}$ reflections | 2173 (145) |
| Final $R_{work}$ (%) | 17.5 (29.1) |
| Final $R_{free}$ (%) | 20.3 (29.5) |
| **No. of non-H atoms** | |
| Protein | 3347 |
| Ligand | 49 |
| Water | 312 |
| Total | 3708 |
| **R.m.s. deviations** | |
| Bonds (Å) | 0.008 |
| Angles (°) | 1.3 |
| **Average B factors** (Å$^2$) | |
| Protein | 42.7 |
| Ligand | 26.6 |
| Water | 44.1 |
| **Ramachandran plot**‡ | |
| Favored regions (%) | 99.0 |
| Additionally allowed (%) | 1.00 |

| | |
|---|---|
| Outliers (%) | 0.00 |

[a] Values in parenthesis are for the highest resolution shell.

[b] Rsym = $\Sigma^{|I-(I)|} / \Sigma^{I}$ where *I* is the observed intensity of a reflection and *(I)* is the average intensity of all the symmetry related reflections.

[c] For $R_{free}$ calculation, 5% randomly selected reflections were excluded from the refinement.

**TABLE S2. Crp-activated and Crp-repressed genes.**

| | Crp-activated[1] | | | | Crp-repressed[2] | |
|---|---|---|---|---|---|---|
| Planktonic | ***ansB*** | *ypeI* | ***ypo1749*** | ***ypo3006*** | *acpS* | *nuoM* |
| | *caf1M* | ***ypeR*** | ***ypo1761*** | *ypo3136* | *bioC* | *nuoN* |
| | ***celA*** | ***YPMT1.70*** | ***ypo1883*** | ***ypo3228*** | *cydC* | ***ptsP*** |
| | ***crp*** | ***ypo0007*** | ***ypo1886*** | ***ypo3287*** | ***cysA*** | ***purE*** |
| | *cspE* | *ypo0176* | ***ypo1887*** | *ypo3318* | ***deaD*** | ***purK*** |
| | *cycA* | ***ypo0277*** | *ypo1904* | ***ypo3472*** | *envZ* | *putA* |
| | *dadA* | ***ypo0315*** | ***ypo1986*** | ***ypo3514a*** | *fhuD* | *recB* |
| | ***fadD*** | ***ypo0402*** | ***ypo1989*** | ***ypo3619*** | ***flgH*** | *recD* |
| | ***fucR*** | *ypo0424* | ***ypo2126*** | ***ypo3647*** | *ftsI* | *recF* |
| | ***glpF*** | ***ypo0628*** | *ypo2127* | *ypo3648* | *ftsL* | *recJ* |
| | ***imm*** | ***ypo0749*** | ***ypo2220*** | ***ypo3681*** | *ftsW* | *rnr* |
| | ***kbl*** | *ypo0760* | *ypo2277* | *ypo3682* | *glgC* | *rpmF* |
| | ***malT*** | ***ypo0856*** | ***ypo2278*** | ***ypo3874*** | *glgP* | *visC* |
| | ***mlc*** | ***ypo0867*** | *ypo2279* | *ypo3995* | ***glyS*** | *wrbA* |
| | ***mtta1*** | ***ypo0884*** | *ypo2337* | *ypo4020* | *hemY* | *ypo0131* |
| | ***ompC2*** | ***ypo0959*** | ***ypo2444*** | ***ypo4036*** | *iucD* | *ypo0147* |
| | ***pim*** | ***ypo0987*** | *ypo2472* | ***ypo4041*** | *livG* | *ypo0381* |
| | ***pla*** | ***ypo1011*** | *ypo2481* | ***YPPCP1.06*** | *mgtE* | *ypo0569a* |
| | ***psaE*** | ***ypo1174*** | ***ypo2589*** | ***YPPCP1.08c*** | *murC* | *ypo1242* |
| | ***rbsK*** | ***ypo1235*** | ***ypo2590*** | ***yspI*** | *murD* | *ypo1243* |
| | ***rcsB*** | ***ypo1255*** | *ypo2729* | | *murE* | *ypo1981* |
| | ***rnk*** | ***ypo1382*** | ***ypo2730*** | | ***murF*** | *ypo2840* |
| | *sfcA* | ***ypo1454*** | ***ypo2863*** | | ***murG*** | *ypo3056* |
| | ***wbyK*** | ***ypo1496*** | ***ypo2955*** | | *nuoE* | *ypo3435* |
| | ***ydjJ*** | *ypo1574* | ***ypo2962*** | | *nuoF* | *ypo3452* |
| | ***ymoA*** | ***ypo1620*** | ***ypo2980*** | | *nuoH* | *ypo3453* |
| | ***yobD*** | ***ypo1718*** | ***ypo3004*** | | *nuoI* | *ypo3523* |
| Biofilm | *aidB* | *YPMT1.69* | *ypo1234* | ***ypo2863*** | *aceA* | *leuB* |
| | ***ansB*** | ***YPMT1.70*** | ***ypo1235*** | *ypo2864* | *cmr* | *leuC* |
| | *argR* | *YPMT1.73* | ***ypo1255*** | *ypo2884* | *cysA* | *luxS* |
| | ***celA*** | ***ypo0007*** | *ypo1341* | ***ypo2955*** | *cysI* | *menC* |
| | ***crp*** | *ypo0099* | *ypo1342* | ***ypo2962*** | *cysJ* | *menE* |
| | ***fadD*** | *ypo0259* | ***ypo1382*** | *ypo2963* | *cysK* | *murF* |
| | *fadH* | *ypo0276* | ***ypo1454*** | ***ypo2980*** | *cysP* | *murG* |
| | *fruB* | ***ypo0277*** | *ypo1463* | *ypo3002* | *deaD* | *ntpA* |
| | ***fucR*** | ***ypo0315*** | *ypo1465* | *ypo3003* | *fabA* | *ompX* |
| | *glpD* | *ypo0339* | *ypo1474* | ***ypo3004*** | *flgG* | *pdxA* |

| | | | | | |
|---|---|---|---|---|---|
| ***glpF*** | ***ypo0402*** | *ypo1477a* | ***ypo3006*** | *flgH* | *proW* |
| *gutB* | *ypo0523* | *ypo1484* | *ypo3137* | *fyuA* | *ptsP* |
| *hexR* | *ypo0536* | *ypo1484a* | *ypo3220* | *glyQ* | *purE* |
| *hpaR* | *ypo0598* | *ypo1492* | ***ypo3228*** | *glyS* | *purK* |
| *hxuB* | *ypo0601* | ***ypo1496*** | ***ypo3287*** | *guaA* | *purL* |
| ***imm*** | ***ypo0628*** | *ypo1498* | *ypo3351* | *guaB* | *recQ* |
| *insA* | *ypo0688* | ***ypo1620*** | ***ypo3472*** | *hemF* | *sodC* |
| ***kbl*** | *ypo0689* | *ypo1651* | ***ypo3514a*** | *hmuU* | *tolQ* |
| *kdgC* | *ypo0690* | *ypo1671* | *ypo3531* | *hscA* | *ybbA* |
| ***malT*** | *ypo0691* | *ypo1687a* | *ypo3611a* | *htpG* | *ybiT* |
| *manX* | ***ypo0749*** | *ypo1707* | ***ypo3619*** | *ileS* | *ygbE* |
| ***mlc*** | *ypo0750* | ***ypo1718*** | ***ypo3647*** | *ilvA* | *yjbA* |
| ***mtta1*** | *ypo0759* | *ypo1739* | ***ypo3681*** | *ilvC* | *ypo2036* |
| *ogl* | *ypo0764* | ***ypo1749*** | *ypo3683* | *ilvD* | *ypo2622* |
| *ogt* | *ypo0768* | ***ypo1761*** | *ypo3709* | *ilvE* | *ypo3033* |
| ***ompC2*** | *ypo0804* | *ypo1811* | *ypo3803* | *ilvM* | *ypo3791* |
| *pckA* | *ypo0806* | ***ypo1883*** | *ypo3839* | | |
| ***pim*** | *ypo0808* | ***ypo1886*** | *ypo3840* | | |
| ***pla*** | *ypo0809* | ***ypo1887*** | ***ypo3874*** | | |
| ***psaE*** | *ypo0813* | ***ypo1986*** | *ypo3885* | | |
| *psaF* | *ypo0814* | ***ypo1989*** | *ypo3893* | | |
| ***rbsK*** | *ypo0833* | *ypo2073* | *ypo3922* | | |
| ***rcsB*** | ***ypo0856*** | ***ypo2126*** | *ypo3923* | | |
| *rhaR* | ***ypo0867*** | *ypo2169* | *ypo3935* | | |
| ***rnk*** | *ypo0874* | ***ypo2220*** | *ypo3953* | | |
| ***wbyK*** | *ypo0881* | ***ypo2278*** | *ypo3981* | | |
| *wbyL* | *ypo0882* | *ypo2282* | *ypo3983* | | |
| *wzz* | *ypo0883* | ***ypo2337*** | *ypo4031* | | |
| *yapA* | ***ypo0884*** | ***ypo2444*** | ***ypo4036*** | | |
| *ybcI* | *ypo0947* | *ypo2482* | *ypo4040* | | |
| ***ydjJ*** | ***ypo0959*** | *ypo2503* | ***ypo4041*** | | |
| *ydjM* | *ypo0965* | *ypo2531a* | *ypo4049* | | |
| *yfiA* | *ypo0978* | ***ypo2589*** | *ypo4050* | | |
| *ylaC* | *ypo0982* | ***ypo2590*** | *ypo4081* | | |
| ***ymoA*** | ***ypo0987*** | *ypo2591* | ***YPPCP1.06*** | | |
| ***yobD*** | ***ypo1011*** | *ypo2611* | ***YPPCP1.08c*** | | |
| ***ypeR*** | *ypo1088* | ***ypo2730*** | *YPPCP1.09c* | | |
| *YPMT1.53c* | ***ypo1174*** | *ypo2733* | ***yspI*** | | |
| *YPMT1.68A* | *ypo1233* | *ypo2778* | | | |

[1]Decreased expression in Δ*crp* vs. *Y. pestis* and glucose vs. glycerol (Log2 FC<1, FDR p<0.05)

[2]Increased expression in Δ*crp* vs. *Y. pestis* and glucose vs. glycerol (Log2 FC>1, FDR $p<0.05$)
**Bolded genes** are Crp-activated or Crp-repressed in planktonic and biofilm states.

Table 1. Gene expression change due to Δ*crp* in metal acquisition genes

| | | Log2 fold-change *Y. pestis*/Δ*crp*[a] | |
|---|---|---|---|
| Group | Gene | Planktonic | Biofilm |
| Feo Iron Transport | *feoA* | +0.509 | **+0.693** |
| | *feoB* | -0.120 | -0.322 |
| Iron Regulation | *fur* | +0.590 | +0.432 |
| Ybt Receptor | *fyuA* | **-2.552** | **-1.764** |
| Heme transport | *hmuR* | **-0.629** | -0.420 |
| | *hmuS* | **-1.111** | -0.611 |
| | *hmuT* | **-1.939** | **-0.925** |
| | *hmuU* | **-2.493** | **-1.067** |
| | *hmuV* | **-1.274** | -0.468 |
| Ybt Synthesis | *irp1* | **-2.575** | **-1.422** |
| | *irp2* | **-2.882** | **-1.880** |
| | *irp3* | **-2.434** | **-1.485** |
| | *irp4* | **-3.701** | **-2.187** |
| | *irp5* | **-3.297** | **-1.912** |
| | *irp6* | **-2.637** | **-2.706** |
| | *irp7* | **-3.168** | **-2.595** |
| | *irp8 (ybtX)* | **-3.870** | **-2.587** |
| Ybt Regulation | *ybtA* | -1.332 | **-2.349** |
| | *ybtS* | **-1.398** | -0.590 |
| Yfe Iron Transport | *yfeA* | +0.320 | +0.552 |
| | *yfeB* | -0.058 | +0.097 |
| | *yfeC* | -0.397 | +0.191 |
| | *yfeD* | +0.036 | -0.179 |
| | *yfeE* | +0.091 | +0.162 |
| | *yfeN* | +0.887 | -0.222 |
| | *yfeY* | +0.476 | +0.252 |
| Zinc Transport | *znuA* | +0.053 | +0.273 |
| | *znuB* | **-1.345** | **-1.249** |
| | *znuC* | -0.741 | -0.425 |

[a]Positive number indicates gene more highly expressed in *Y. pestis*/Δcrp and thus is Crp-activated. Negative number indicates gene indicates gene is Crp-repressed.

Bold represents FDR $p$ <0.05

Table 2. Gene expression change due to Δ*crp* in quorum sensing genes

| Group | Gene | Log2 fold-change *Y. pestis*/Δ*crp*[a] Planktonic | Biofilm |
|---|---|---|---|
| AHL | *ypeI* | **+1.817** | **+1.555** |
| | *ypeR* | **+1.624** | **+1.446** |
| | *yspI* | **+2.792** | **+2.709** |
| | *yspR* | **+2.731** | **+2.189** |
| AI-2 | *luxS* | -0.665 | **-1.099** |

[a]Positive number indicates gene is Crp-activated. Negative number indicates gene is Crp-repressed.

Bold represents FDR $p < 0.05$

**TABLE S3** Bacterial Strains and Plasmids used in this study

| Strain or plasmid | Relevant description | Source or Reference |
|---|---|---|
| ***Yersinia pestis* CO92** | | |
| SAN3 | pCD1+, *pgm+,* pMT1+ | (74) |
| PAN30 | pCD1-, Δ*cyaA* | This study |
| PAN259 | pCD1-, "wild-type" | (74) |
| PAN933 | pCD1-, Δ*crp* | (6) |
| PAN977 | pCD1-, Δ*ypeIR* | This study |
| PAN978 | pCD1-, Δ*yspIR* | This study |
| PAN981 | pCD1-, Δ*luxS* | This study |
| PAN985 | pCD1-, Δ*ypeIR*Δ*yspIR* | This study |
| PAN1071 | pCD1-, Δ*malT* | This study |
| PAN1072 | pCD1-, Δ*crp* + *crp* | This study |
| PAN1073 | pCD1-, Δ*malT* + *malT* | This study |
| ***Rhizobium radiobacter*** | | |
| PAN1045 | pZLR4 | ATCC® BAA-2240 |
| ***Vibrio harveyi*** | | |
| PAN980 | MM32 | (75) |
| ***Escherichia coli*** | | |
| CC118 λ*pir* | *Δ(ara-leu) araD ΔlacX74 galE galK phoA20 thi-1 rpsE rpoB argE(Am) recAl λpir* | (72) |
| DH5α | *F⁻,*φ80d*lacZ*ΔM15 Δ*(lacZYA-argF)*U169 *deoR recA1 endA1 hsdR17(*rK$^-$, mK$^+$*) phoA supE44 λ$^-$ thi-1* | Laboratory stock |
| DH5α λ*pir* | As above with λ*pir* | Laboratory stock |
| S17-1 λ*pir* | Tp$^R$ Sm$^R$ *recA*, *thi-1*, *pro*, *hsdR*$^-$ M+RP4: 2-*Tc*:*Mu*: Km Tn*7* λpir | Laboratory stock |
| TOP10 | F$^-$ *mcrA Δ(mrr-hsdRMS-mcrBC)* φ80*lacZΔ*M15 *ΔlacX74 recA1 araD139* | Invitrogen |

| | | |
|---|---|---|
| | *Δ(ara-leu)7697 galU galK* $\lambda^-$ *rpsL(Str*$^R$*) endA1 nupG* | |
| BL21(DE3)(Magic) | Overexpression of proteins expressed under T7 promoter with plasmid expressing rare codons, Km$^R$ | A. Joachimiak, Argonne National Labs |
| **Plasmids** | | |
| pWL213 | Constitutive expression of *gfp,* Km$^R$ | (35) |
| pUC18 | pUC18R6K-mini-Tn7t, Km$^R$ | (58) |
| pTNS2 | Conjugative helper plasmid Amp$^R$ | (58) |
| pKD13 | Source of pSkippy Kan cassette Km$^R$ | Laboratory Stock |
| pSkippy | IPTG-inducible FLP recombinase, Amp$^R$ | Laboratory Stock |
| pWL204 | Lambda red recombinase, Amp$^R$ | Laboratory Stock |
| pMCSG53 | Vector for protein overexpression Amp$^R$ | (76) |
| pIDP97063 | Overexpression of 6xHis-tagged CRP | This study |
| pLB30 | Promoterless *gfp* for insertion into *Y. pestis* Tn*7* *att* site, Km$^R$ | (35) |
| pJR32 | P*ypeI-gfp,* Amp$^R$ | This study |
| pJR33 | P*yspI-gfp,* Amp$^R$ | This study |
| pJR34 | P*yspR-gfp,* Amp$^R$ | This study |
| pJR35 | P*ypeR-gfp,* Amp$^R$ | This study |
| pJR36 | *crp* complementation, Amp$^R$ | This study |
| pJR37 | *malT* complementation, Amp$^R$ | This study |

**TABLE S4** Oligonucleotides used in this study

| Designation | Sequence (5’ to 3’) | Purpose |
|---|---|---|
| 5’ p1 for pKD13 | GTGTAGGCTGGAGCTGCTTC | Amplification Kan cassette for lambda red |
| 3’ p4 for pKD13 | ATTCCGGGGATCCGTCGACC | Amplification Kan cassette for lambda red |
| 5’ -500 malt | TGCTCGGTGCGGGCTGAGCAG | Lambda red deletion of *malt* |
| 3’ +1 malT P1 | GAAGCAGCTCCAGCCTACACTCTGT GTCCGCTGGAAAAG | Lambda red deletion of *malt* |
| 5’ malt p4 | GGTCGACGGATCCCCGGAATCGGG CGCACACGTGTTTGATG | Lambda red deletion of *malt* |
| 3’ +500 malt | TGCATGACCGGCTTCGTAAC | Lambda red deletion of *malt* |
| 5’ -500 ypeIR | CACTCCCCCAAAGTAGACAAC | Lambda red deletion of *ypeIR* |
| 3' P1 ypeIR | GAAGAGCTCCAGCCTACACATTTAT TTAATTAAACCAATATC | Lambda red deletion of *ypeIR* |
| 5' P4 ypeIR | GGTCGACGGATCCCCGGAATAGCA GACCAAATTTACTTTATCC | Lambda red deletion of *ypeIR* |
| 3' +500 ypeIR | CGGCTGTGCTTTAGCAGCCC | Lambda red deletion of *ypeIR* |
| 5' -500 yspIR | TCCTCCAGCGCTTTGAGACAC | Lambda red deletion of *yspIR* |
| 3' P1 yspIR | GAAGCAGCTCCAGCCTACACTATCT TTCCTGATATTTAATAC | Lambda red deletion of *yspIR* |
| 5' P4 yspIR | GGTCGACGGATCCCCGGAATTCCCT TTCTCCATTTACTGTAC | Lambda red deletion of *yspIR* |
| 3’ +500 yspIR | AGAATTTATGCGATTCGTGGC | Lambda red deletion of *yspIR* |
| 5’ SpeI -500 crp | TTCGATCATGCATGAGCTCAAAAAT AGACACGACATCAATG | Tn7 complementation of *crp* |
| 3' SpeI +500 crp | CCTGCAGCCCGGGGGATCCACCAT GCTGAGACTGAAAATAG | Tn7 complementation of *crp* |
| 5’ SpeI -500 malT | TTCGATCATGCATGAGCTCATGCTC GGTGCGGGCTGAGCAG | Tn7 complementation of malt |
| 3’ SpeI +500 malt | CCTGCAGCCCGGGGGATCCATGCA TGACCGGCTTCGTAAC | Tn7 complementation of malt |
| 5' -300 pla BT | 5BIOSG/TCTCGCCCGTAAATACCTG AG | Biotinylated primer for amplifying *pla* promoter |
| 3' +1 pla BT | 5BIOSG/TAGACACCCTTAATCTCTCT G | Biotinylated primer for amplifying *pla* promoter |
| 5’ -300 ptsG BT | /5Biosg/TTA GCT CGT AAT TAA TCA CCG | Biotinylated primer for amplifying *ptsG* promoter |

| | | |
|---|---|---|
| 3' +1 ptsG BT | /5Biosg/ATA GTT GAG CGT GCT CCT GAG | Biotinylated primer for amplifying *ptsG* promoter |
| 5' -300 malT BT | 5BIOSG/GTGAATGGGTAATTTTACCCG | Biotinylated primer for amplifying *malT* promoter |
| 3' +1 malT BT | 5BIOSG/ATTCTGTGTCCGCTGGAAAAG | Biotinylated primer for amplifying *malT* promoter |
| 5' -300 ypeR BT | 5BIOSG/ATATATTCACAGCAGACTAAC | Biotinylated primer for amplifying *ypeR* promoter |
| 3' +1 ypeR BT | 5BIOSG/AGCAGACCAAATTTACTTTATC | Biotinylated primer for amplifying *ypeR* promoter |
| 5' -300 yspI BT | /5BIOSG/CATGCTACGAGAGAATTATAC | Biotinylated primer for amplifying *yspI* promoter |
| 3' +1 yspI BT | 5BIOSG/TATCTTTCCTGATATTTAATAC | Biotinylated primer for amplifying *yspI* promoter |
| 5’ -300 yspR BT | /5Biosg/GAG GGA CAA TTA ATC ACA ATG | Biotinylated primer for amplifying *yspR* promoter |
| 3' +1 yspR BT | /5Biosg/TCC CTT TCT CCA TTT ACT GTA C | Biotinylated primer for amplifying *yspR* promoter |
| 5' -300 ypeI BT | /5Biosg/CTA AAT ACC ACT CAC ACA AGC | Biotinylated primer for amplifying *ypeI* promoter |
| 3' +1 ypeI BT | /5Biosg/ATT TAT TTA ATT AAA CCA ATA TC | Biotinylated primer for amplifying *ypeI* promoter |
| gyrB.r.3' | ATTGGTAAAGGTCTGGAAACTTGGCC | qRT-PCR of *gyrB* |
| gyrB.f.5' | TCGCCGTGAAGGTAAAGTTC | qRT-PCR of *gyrB* |
| 5’ pla 397 | GACCTCAATGTGAAAGGCTGGTTACACC | qRT-PCR of *pla* |
| 3’ pla 498 | ACCACCTGTAGCTGTCCAACTGAAAC | qRT-PCR of *pla* |
| 5’ ptsG 132 | CTC TCA CGT AAT GGC AGA AG | qRT-PCR of *ptsG* |
| 3’ ptsG 229 | ACC GTC GTT GTT GGT AAA G | qRT-PCR of *ptsG* |
| 5' malt1078 | GCTATCCACCACGCTTTAG | qRT-PCR of *malT* |
| 3' malt 1184 | GTTCCAACAACGCCAATTC | qRT-PCR of *malT* |
| 5’ ybta 282 | TCGCCATTACATTACCCATAC | qRT-PCR of *ybtA* |
| 3’ ybta 414 | CAGTGGGAGTCGATCTTATTC | qRT-PCR of *ybtA* |
| 5’ fyua 848 | AGACCCTGAGTGGGAAATAC | qRT-PCR of *fyuA* |
| 3’ fyua 1027 | GTACAGCCCAAACACCATATC | qRT-PCR of *fyuA* |
| 5’ yspI 27 | CGACGAGTTGACCGATATAC | qRT-PCR of *yspI* |
| 3’ yspI 187 | ACGCACACTGCAGATTAG | qRT-PCR of *yspI* |
| 5’ yper 281 | ATTCGGCCGTGTTCAATC | qRT-PCR of *ypeR* |
| 3’ yper 412 | TGTTCACCTCGATGCTTTC | qRT-PCR of *ypeR* |
| 5’ crp 2 | TGGTTCTCGGTAAGCCACAAACAGAC | qRT-PCR of *crp* |
| 3’ crp 146 | GCAACGGAGCCTTTCACGATGTAG | qRT-PCR of *crp* |
| 5' gfp +1 | ATGAGTAAAGGAGAAGAACTTTTC | Amplification of *gfp* CDS |

| | | |
|---|---|---|
| 3' gfp SpeI puc18 | CAGCCCGGGGGATCCACTAGTTATT TGTATAGTTCATCCATG | Amplification of *gfp* CDS for Tn7 |
| 5’ puc18 pypei | TCATGCATGAGCTCACTAGCACTCC CCCAAAGTAGACAAC | Amplification of P*ypeI*-GFP reporter |
| 3’ gfp pypei | AGTTCTTCTCCTTTACTCATAAATA CTTTTAACATAATAAAAAC | Amplification of P*ypeI*-GFP reporter |
| 5’ puc18 pyper | TCATGCATGAGCTCACTAGCGGCTG TGCTTTAGCAGCCC | Amplification of P*ypeR*-GFP reporter |
| 3’ gfp pyper | AGTTCTTCTCCTTTACTCATTTCATT ATCAAAAAAATTAATTATC | Amplification of P*ypeR*-GFP reporter |
| 5’ puc18 pyspi | TCATGCATGAGCTCACTAGTCCTCC AGCGCTTTGAGACAC | Amplification of P*yspI*-GFP reporter |
| 3’ gfp pyspi | AGTTCTTCTCCTTTACTCATGTATCT GACATCGAAAATTTC | Amplification of P*yspI*-GFP reporter |
| 5’ puc18 pyspr | TCATGCATGAGCTCACTAGGGCTGT AAGAATTTATGCGATTC | Amplification of P*yspR*-GFP reporter |
| 3’ gfp pyspr | AGTTCTTCTCCTTTACTCATGTTACT CCTATTGAAAACAGAATG | Amplification of P*yspR*-GFP reporter |

**Dataset S1 1.0 MB (.xlsx) Uploaded as separate file**

**Dataset S2 188 kB (.xlsx) Uploaded as separate file**

**Dataset S3 37 kB (.xlsx) Uploaded as separate file**
